# DEK-rRNA interactions regulate ribosome biogenesis and stress response

**DOI:** 10.1101/2024.08.23.609341

**Authors:** Nengwei Xu, Kunqi Chen, Malte Prell, Pengtao Liang, Shuhuai Yang, Shengjie Guo, Yuli Lu, Soham Dorle, Samia Idaghdour, Ferdinand Kappes

**Affiliations:** Department of Biological Sciences, School of Science, Xi’an Jiaotong-Liverpool University, Suzhou, Jiangsu, China; Institute for Biochemistry and Molecular Biology, Medical School, RWTH Aachen University, 52074 Aachen, Germany; Division of Natural and Applied Sciences, Duke Kunshan University, Kunshan, Jiangsu, China

## Abstract

DNA/chromatin-based functions of the DEK oncogene, a unique nucleic acid-interacting factor in metazoans, have been widely investigated, yet its role in cellular RNA biology is under-studied. Herein we employed CLIP-seq alongside mutational, biochemical, and cellular approaches to gain deeper insights into the cellular DEK-RNA interplay. We confirm interaction of DEK with coding RNA, yet also report association with ribosomal RNA (rRNA) and thereby establishing a robust link between DEK and ribosome biology. Indeed, cells lacking DEK showed marked deficits in cytoplasmic ribosome quality and function. This phenotype was exclusively rescued by C-terminal DEK, harboring two RNA interaction domains, but not by an rRNA-binding deficient mutant. Mechanistically, we uncovered pleiotropic involvement of DEK in RNA polymerase I-mediated rRNA transcription and processing pathways. More specifically, we found direct interaction of DEK with RNA polymerase III-transcribed 5S rRNA and identified DEK as a regulator of the Impaired Ribosome Biogenesis Checkpoint (IRBC). Within this ribosomal stress pathway, DEK depletion results in free 5S RNP, triggering stabilization of p53 via inhibition of MDM2. In summary, our multilayer analysis revealed DEK as a potent cellular RNA binding protein and provides first evidence of DEK as a regulator of ribosome biogenesis and stress response via the 5S RNP-MDM2-p53 axis.

## Introduction

Since its isolation from the DEK-NUP214 fusion gene in acute myeloid leukemia ^1^ over three decades ago, research on the biochemically distinct and multifunctional chromatin-associated human DEK protein has consistently revealed tight associations between its functions and disease pathogenesis, particularly pertinent to cancer and autoimmune diseases. A growing number of studies have identified overexpression of DEK in an array of solid tumors ^2^, proposed DEK as a potential serum and urinary cancer biomarker due to its non-classical secretion ^3–6^ and have established DEK as a *bona fide* oncogene ^7–9^. Even though the precise molecular details underlying its pathogenicity and tumorigenicity remain largely obscure, DEK functions contribute to tumorigenesis and cancer maintenance in a variety of specific, yet also pleiotropic ways ^10, 11^. DEK is involved in signaling pathways and oncogenic activities, including p53-dependent/independent cellular senescence ^8, 12–15^, regulation of apoptosis ^16^, and cell proliferation through PI3K-Akt-mTOR ^17^ and Rho signaling ^18^. Biochemical and structural analyses uncovered DEK as a predominantly nuclear protein under steady-state conditions ^4, 5, 19–24^, whose 375 amino acids contain a unique pseudo-SAP/SAP DNA interaction domain ^25^. This domain (DEK 87-187) can introduce energy-independent supercoils to DNA, and is, in concert with the C-terminal DNA binding and phosphorylation-dependent multimerization domain (DEK 250-375), responsible for its perceived principal molecular function in chromatin organization and maintenance ^26–30^. Such interactions of DEK with cellular DNA are predominantly based on structure rather than sequence, and are considered foundational to DEK’s pleotropic functions in a range of nuclear processes, including DNA replication ^31–33^, DNA damage repair ^5, 34–36^, and transcriptional regulation ^37, 38^.

However, interactions of DEK with cellular nucleic acids are not limited to DNA as very early studies found DEK associated with ribonucleic particles and as a factor involved in mRNA splicing ^39–44^. Concordant with this, our own early observation showed that ∼ 10% of a given DEK population is associated with cellular RNA ^19^. In full support of these earlier reports, the RBP2GO server (https://rbp2go.dkfz.de/) now lists 14 out of 43 recent unbiased global RNA interactome capture studies as independently identifying DEK as a cellular mRNA binding protein (RBP) (**Supplementary File 1**). Using both poly(A) and non-poly(A)-based approaches, cellular DEK-mRNA interactions were found in HEK293, HuH7, HeLa, MCF10A, U2-OS and embryonic stem cells ^45–47^. Additionally, three studies mapped amino acids 332-344 ^47, 48^ or 336-349 ^49^ in DEK as pivotal for interaction with mRNA and thus suggesting dual-functionality of the C-terminal DNA binding domain of DEK, spanning aa 250 to 375. In contrast, no evidence of mRNA binding of DEK was found in studies employing Serial RNA interactome capture (serlC), photo/peptide-crosslinking and affinity purification (pCLAP) in K562, HeLa or HEK293T cells, respectively ^50–52^. However, collectively the amassed data clearly suggests general roles for DEK in cellular (m)RNA biology, which remains an under-studied aspect of DEK biology^45–47, 49, 53^.

Besides interaction with coding RNA, a genome-wide screen in human cells recognized DEK as a ribosomal RNA (rRNA) processing factor ^54^. Additionally, systematic proteomic analysis revealed the presence of DEK in human nucleoli, which is the location where ribosomal RNA genes are clustered within the nucleus and thus signifies the starting point of ribosome biogenesis ^55^. The recently identified DEK and DEK-NUP214 interactomes further solidify associations of DEK with ribosome biology, as exemplified, amongst others, by interaction of DEK with a number of proteins derived from the large ribosomal subunit (e.g. RPL11 and RPL5) ^56, 57^. However, how DEK participates in ribosome biology remains elusive.

Ribosome biogenesis is an essential, highly complex, extremely energy-consuming process with a multitude of dynamically coordinated steps that require concerted action of all three RNA polymerases (pol I, II, III) ^58^. It is initiated in the nucleolus by pol I-dependent transcription and regulated via dynamic co-transcriptional processing of a polycistronic pre-47S rRNA. This is followed by export of pre-mature ribosomal subunits, composed of ribosomal proteins and rRNAs to eventually form translation-competent ribosomes in the cytoplasm of human cells, which, in addition, requires the assistance of more than 200 other ribosomal factors ^59^. As cell proliferation is intimately tied to the rate of ribosome biosynthesis and ribosome function, alterations to ribosome homeostasis and/or quality are considered critical steps in cancer pathogenesis ^60, 61^. Not surprisingly, nucleolar/ ribosomal stress typically triggers activation of crucial stress pathways, such as the Impaired Ribosome Biogenesis Checkpoint (IRBC). Within this pathway, mis-localized, non-ribosome bound 5S RNP, comprised of proteins RPL5, RPL11 and 5S rRNA, results in attenuation of the MDM2 ubiquitin ligase activity and thus stabilization of p53, highlighting key functions for p53 in sensing ribosomal dysfunction ^62, 63^.

In addition to the unbiased RNA interactome capture studies noted above, which identified the C-terminal DNA binding domain DEK 250-375 as an RNA interaction domain, we have very recently independently shown that DEK is indeed an RNA interacting factor *in vitro* and in cells using a newly developed bacterial screening technique (BGIS). Specifically, we have mapped DEK 187-270, an intrinsically disordered domain, as an RNA interaction domain in DEK ^64^. Thus, it appears that RNA interactions of DEK are mediated predominantly via its C-terminal portion DEK 187-375, involving an intrinsically disordered domain and a dual-specificity nucleic acid interaction domain that can bridge contacts to DNA and RNA molecules. Yet, as little is known about the precise molecular details governing DEK-RNA interactions, we applied RNA crosslink immunoprecipitation followed by high throughput sequencing (CLIP-seq) and a range of other biochemical and cellular assays to decipher such interactions and their functional consequences on the cellular level in this study.

## Results

### Bacterial Growth Inhibition Screen (BGIS) selects a DEK 187-270 mutant with attenuated RNA binding ability

Having recently identified an RNA-interacting region in the DEK protein localizing to amino acids 187-270 ^64^, we aimed to first create mutants in this region with attenuated RNA binding ability to perform more detailed structure-function studies in cells. However, a targeted mutational approach proved difficult due to the following reasons: i) DEK 187-270 contains 25 positively charged lysine and arginine residues (approximately 30% of this fragment) (**Supplementary Figure 1A**); ii) NMR data ^65,66^ and structure prediction servers (NORSnet, DISOPRED2, PROFbval and Ucon) ^67–69^ identify this region as intrinsically disordered (**Supplementary Figure 1B** and C), and iii) no prominent disease-related mutational hotspots were identified (**Supplementary Figure 1D, Supplementary File 2**). As this information established no clear outline for a mutational strategy, we employed a recently developed approach, Bacterial Growth Inhibition Screen (BGIS) ^27, 64^, in combination with random mutagenesis, to streamline selection of RNA-binding deficient mutants in DEK 187-270. This method yielded three suitable GST-tagged mutants, which were expressed and purified from *E. coli* (**Figure 1A and B; Supplementary Figure 2)**. Even though RNA-EMSA indicated only modestly reduced binding affinity to single stranded RNA for each of these mutants individually (**Figure 1C; Supplementary Figure 3**), a combination of all mutations from clones RBN#36, #267 and #696 into one mutant, now termed “RBN#3c” (three mutants combined), clearly resulted in a marked reduction of RNA binding affinity within the DEK 187-270 fragment (**Figure 1C**). To employ this mutant for subsequent cellular assays, stable and inducible cell lines were established via lentiviral delivery. These cells showed robust and comparable expression of eGFP-DEK fusions, including full length DEK containing all mutations found in RBN#3c (**Supplementary Figure 4**) ^70^.

**Figure 1.**
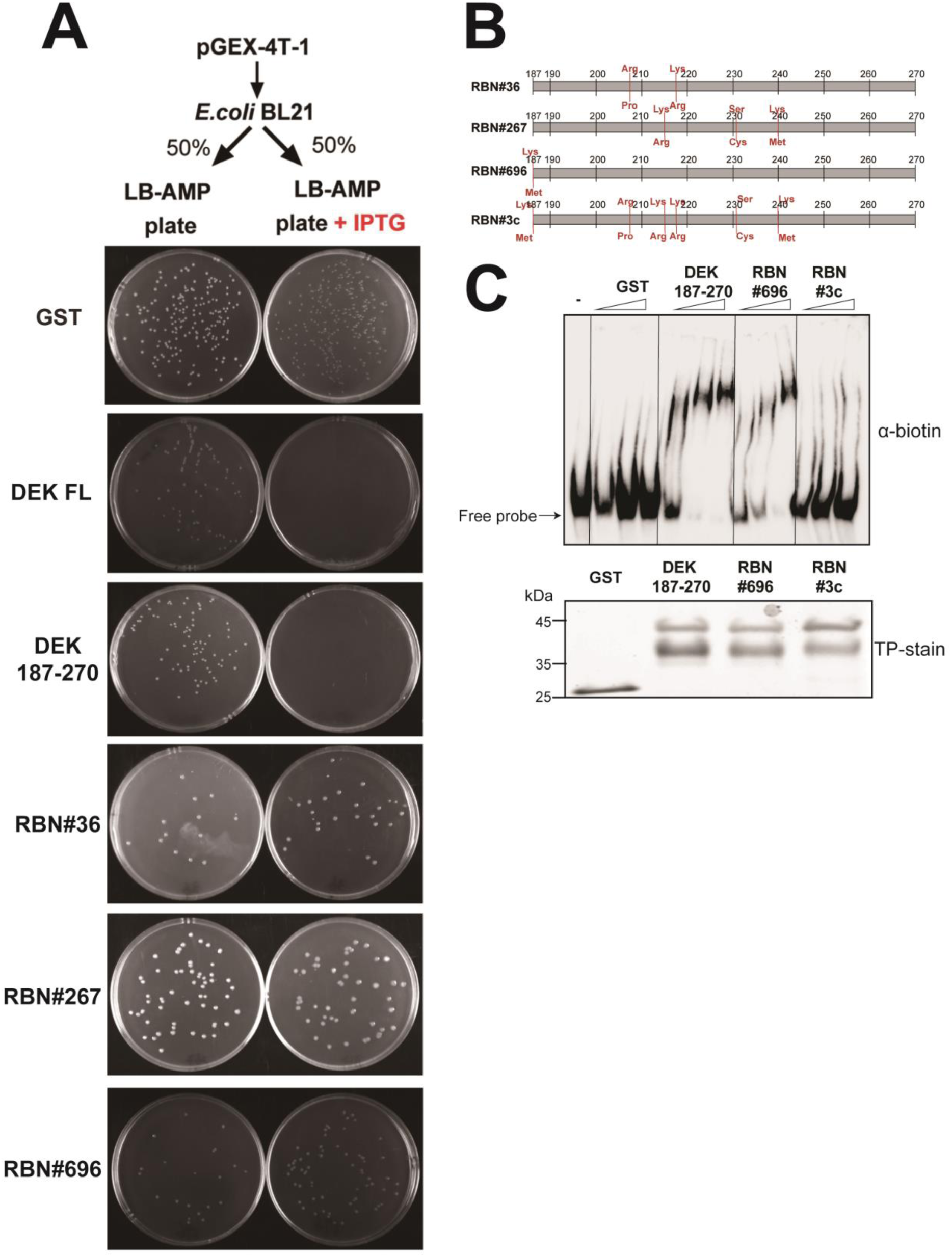
Screen for a DEK mutant attenuated in its ability to bind RNA via random mutagenesis and Bacterial Growth Inhibition Screen (BGIS). **A.** pGEX-4T-1 plasmids containing wild-type full length DEK, the fragment 187-270 and three mutants obtained via BGIS (Supplementary Figures 2 and 3) were transformed to *E. Coli* BL21 and plated as described on top. Micrographs of plates were taken the next day indicating growth inhibition for DEK full length and the DEK187-250 fragment, yet not GST alone or the selected mutants; B. Schematic depiction of identified mutations of positive hits from (A). Original amino acids are denoted on top and resulting changes on the bottom; C. RNA-EMSA: 10 ng of RNA pentaprobe was incubated with increasing concentrations (0.3, 0.6 and 1.2 pmol) of purified recombinant protein (GST alone or with indicated GST-DEK187-270 fusions) or left untreated (-). RNA binding activity was subsequently analyzed by native polyacrylamide gel electrophoresis followed by detection of biotinylated RNA via chemiluminescence. Purified proteins used for each reaction were analyzed by SDS-PAGE and Coomassie Blue staining (TP-stain) confirming equimolar input to RNA-EMSA reactions.

### CLIP-seq reveals interaction of ectopically expressed DEK with specific sets of coding and non-coding RNA

Using this inducible cell system we next performed UV crosslinking and immunoprecipitation followed by sequencing (CLIP-seq) to harness cellular DEK-interacting RNAs with high stringency and in comparison, between wt full length DEK (aa 1-375) and the RNA-binding RBN#3c full length mutant (aa 1-375). After systematic optimization of the CLIP conditions (**Supplementary Figure 5**), eGFP-DEK or eGFP expressed in DEK KO HeLa S3 cells was captured by GFP trap purification. After on-bead biotinylation of resulting ribonucleoprotein complexes and separation on polyacrylamide gels, RNA was detected via streptavidin antibodies (**Figure 2A**). Whereas eGFP alone produced negligible signals, substantial signals above 70 kDa, the proper molecular weight of eGFP-DEK, can be seen in the eGPF-DEK CLIP sample. This indicated successful capture of crosslinked RNA molecules to DEK and thus broadly confirmed DEK as a cellular RNA binding factor. Ribonucleoprotein complexes corresponding to RNA smaller than 200 nt (∼70 to 130 kDa) were extracted from two independent CLIP samples for each condition (DEK wt_1, DEK wt_2 and RBN#3c_1, RBN#3c_2) and the resulting purified RNA was subjected to adaptor ligation, cDNA library construction and sequencing. Overall, more than 60 million clean reads were generated from both DEK wild-type (wt) CLIP samples individually (DEK wt_1 and DEK wt_2). The RBN#3c mutant also yielded substantial amounts of total RNA reads, yet fewer than those from DEK wt, with approximately 47 million and 46.7 million reads, respectively (**Figure 2B**). As this was surprising, we tested DEK 250-375, a known DNA interaction domain in DEK, in RNA-EMSA, which we did not assess in isolation in our previous study ^64^. This indeed also confirmed this C-terminal portion of DEK as a RNA interacting region (**Supplementary Figure 6**) and further confirming amino acids 332-344 ^53^ or 336-349 ^49^ as mRNA interaction points.

**Figure 2.**
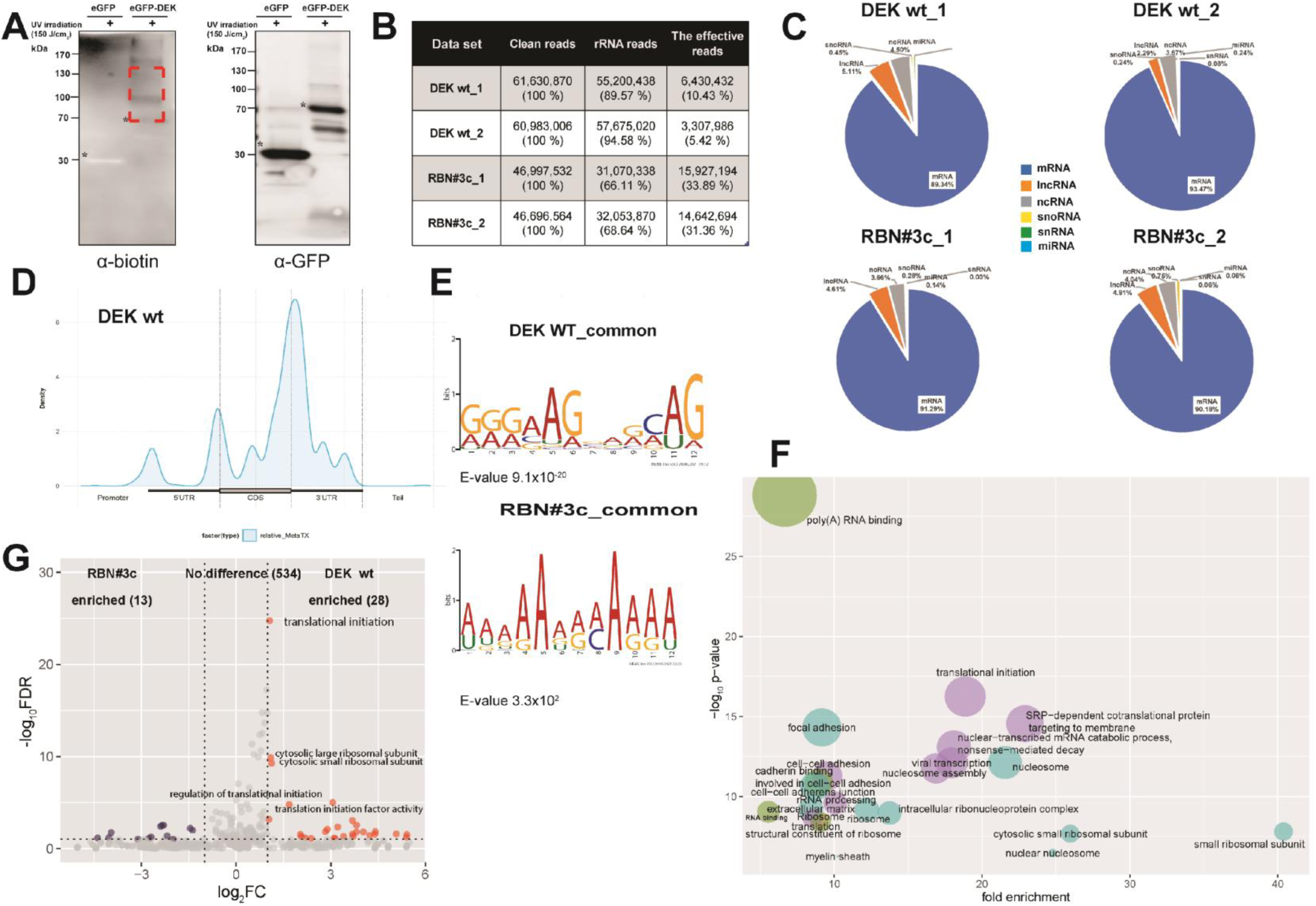
CLIP-seq from HeLa cells identifies a set of coding and non-coding RNA interacting with ectopically expressed DEK. A. Crosslinked ribonucleoprotein complexes from immunoprecipitations of eGFP alone or eGFP-DEK wt in DEK KO HeLa cells were subjected to SDS-PAGE and probed with the antibodies specific to biotin (left panel) or GFP (right panel). Asterisks indicate the positions of eGFP tagged protein immunoprecipitated by GFP-trap. A region corresponding to approximately 50 kDa above the target protein as indicated by the red dashed box was excised for CLIP-seq library preparation; **B.** Table summarizing the number of reads uniquely mapped to the human genome (clean reads) for CLIP-seq biological replicates of both DEK wildtype (DEK wt_1 and DEK wt_2) and mutant (RBN#3c_1 and RBN#3c_2). The percentage and quantity of ribosomal RNA and non-ribosomal RNA (the effective reads) for each sample is indicated; **C.** Pie charts depicting the percentage of CLIP reads from both DEK wt and RNB3#c duplicates mapped to any type of non-ribosomal RNA. In both DEK wt biological replicates, mRNA represents the largest fraction cross-linked to DEK, with 89.34% and 93.47%, respectively, followed by various types of non-coding RNAs. mRNA: messenger RNA, lncRNA: long non-coding RNA; ncRNA: non-coding RNA, snoRNA: small nucleolar RNA, snRNA: small non-coding RNA and miRNA: microRNA; **D.** Distribution of protein-coding transcripts cross-linked to DEK wt mapped to mRNA as analyzed by Guitar R package; **E**. MEME motif searching algorithm identified a purine-rich reoccurring sequence pattern analyzed from the common reads of DEK wt, whereas no statistically significant consensus sequence pattern was found from RBN#3c CLIP samples. **F**. Bubble chart depicting functional clustering of DEK wt interacting mRNA transcripts contained in the functional annotation clusters identified by DAVID bioinformatic tool with a threshold of -log_10_p-value lower than 30. **G.** The fold enrichment value of each annotation cluster derived from DEK wt CLIP datasets generated by DAVID bioinformatics tool was normalized to the one from RBN#3c, giving rise to the red dots indicating enriched GO terms in DEK wt, while blue ones are over-represented in RBN#3c dataset.

After normalization of non-rRNA reads to matched input RNA, we analyzed common calling peaks between the two samples via mapping to the human genome (hg19). This revealed predominant binding of DEK to protein-coding mRNAs (∼90%), followed by several rather small subsets of non-coding RNAs, including lncRNA, ncRNA, snoRNA, snRNA and miRNA (**Figure 2C**, see **Supplementary File 3** for all CLIP’ed RNA species). Besides a quantitative decrease in rRNA reads, general binding preferences of RBN#3c to cellular RNA species appeared initially virtually unchanged (**Figure 2C**). Analysis of the binding distribution of DEK wt on captured mRNA revealed preferential binding to exons, specifically at the 3’ end of exon sequences in proximity to the 3’UTR (**Figure 2D**). Moreover, the motif search algorithm MEME returned a purine-rich reoccurring sequence in DEK wt when using a search window of 8-12 nucleotides. This, however, was altered with RBN#3c, suggesting qualitative differences in the mRNAs captured (**Figure 2E**), which may indicate intricate cooperativity and mutual regulation between DEK domains 187-250 and 250-375 in bridging mRNA contacts.

We next used Gene ontology (GO) analysis to functionally annotate the CLIP’ed mRNAs. This returned overrepresentation of GO terms relating to ribosome biology, including “translational initiation” and “ribosomal subunit” (**Figure 2F**). Interestingly, comparative enrichment analysis between DEK wt and RBN#3c CLIP datasets showed substantial qualitative differences between these data sets. Most strikingly, a loss of ribosome-centric GO terms for RBN#3c samples was observed (**Figure 2H**, see **Supplementary Tables 1 and 2** for GO terms). Taken together, our findings thus far pinpoint at roles for DEK in ribosome biology.

### DEK 187-375, but not RBN#3c, rescues deficiencies in protein translation in DEK depleted cells

To test a potential role of DEK in ribosome function, we applied SUrface SEnsing of Translation (SUnSET) assays ^71^. Via competitive incorporation of puromycin, a tyrosyl-tRNA analogue, relative changes to new cellular protein synthesis can be monitored in immunoblots of total cell lysates via puromycin-specific antibodies. Indeed, substantially reduced protein translation was observed in HeLa S3 (**Figure 3A, lane 1, 2 and 4**), HEK293, and primary human dermal fibroblast (HDF) upon knockdown or knockout of DEK (**Supplementary Figure 7**). Classic ^35^S methionine incorporation experiments with wildtype and DEK KO HeLa cells further supported this observation (**Supplementary Figure 8**). To test whether this marked effect on protein translation is caused exclusively by DEK depletion or by other side effects, rescue experiments with a set of epitope-tagged DEK fusions were carried out. Remarkably, re-expression of FLAG-MYC-DEK (**Figure 3A, lane 6**) or HA-FLAG-DEK (**Figure 3A, lane 7**) in DEK KO cells and eGFP-DEK, as shRNA escape version in DEK KD cells, largely restored the protein synthesis rate in DEK-deficient cells (**Figure 3A, lane 3**). Furthermore, this result eliminated concerns regarding the molecular functionality of the DEK fusions utilized in CLIP.

**Figure 3.**
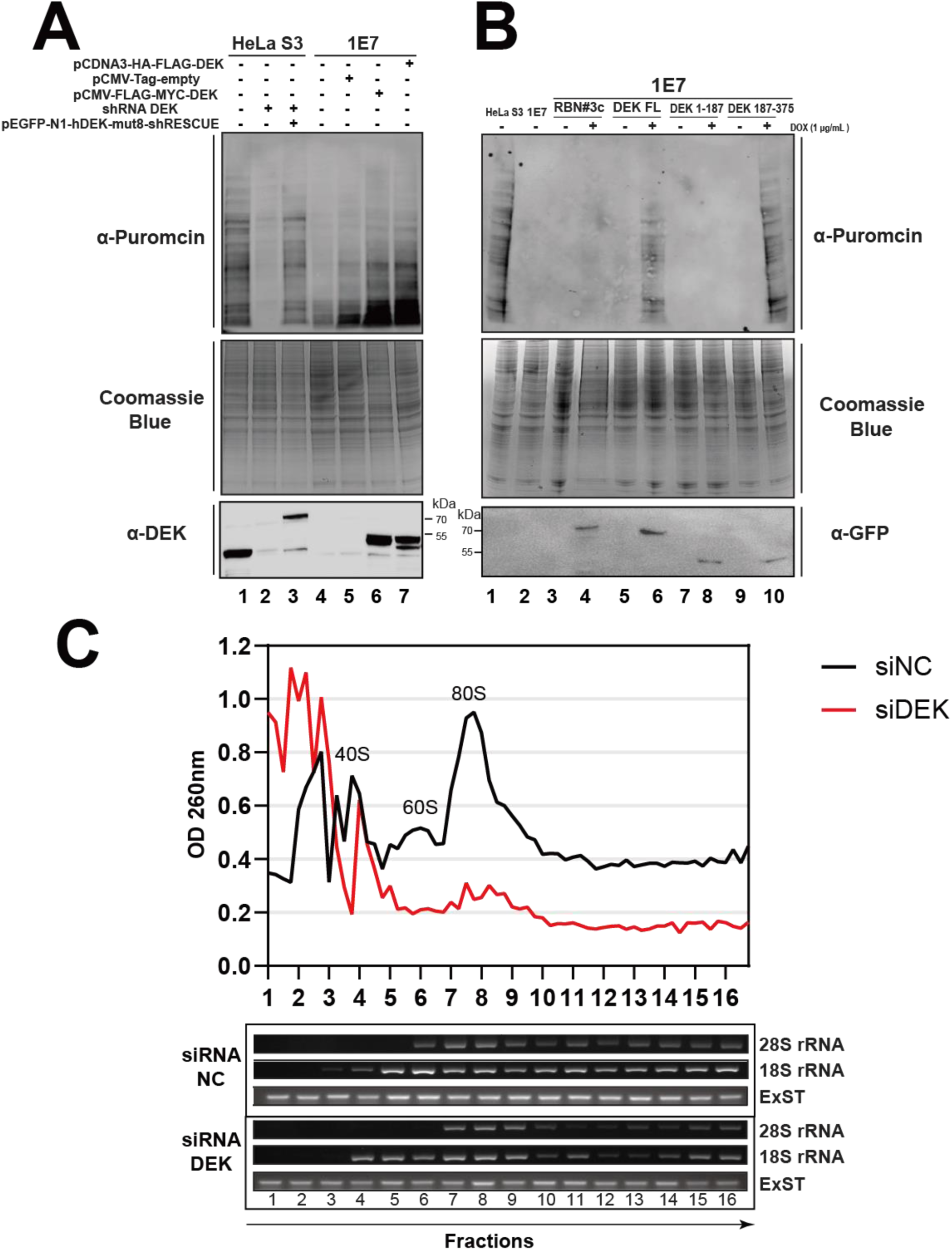
Defective ribosome biogenesis upon DEK depletion results in reduced protein translation. A. SUrface SEnsing of Translation (SUnSET) assay. HeLa S3 wt, DEK KO (1E7) or DEK KD (shRNA) cells transfected with indicated plasmids were incubated with puromycin at a final concentration of 10 μg/mL followed by total protein extraction. Half of the resulting extracts was either separated by SDS-PAGE and analyzed by immunoblotting with antibodies specific to puromycin (12D10) (top) or DEK (bottom), or by SDS-PAGE and Coomassie Blue staining to monitor total protein amounts (middle); **B.** SUnSET assay as in A with DEK KO cells expressing indicated eGFP-DEK fusions after induction with doxycycline. GFP-specific antibodies were used for monitoring expression of respective DEK fragment; Shown is one representative example (see Supplementary Figure 9 for repetitions). **C.** Cytoplasmic ribosome profiling in HeLa S3 cells transfected with siRNAs targeting DEK or with nonsense siRNAs. Each fraction was measured by OD_260_, recorded, and plotted. An aliquot of each fraction was purified and analyzed by RT-qPCR with primers designed for 18S and 28S rRNA to identify the positions of 40S, 60S and 80S ribosomes. ExST: External Standard. Shown is one representative experiment out of two (see also **Supplementary Figure 10** and 11).

To next elucidate the specific domain(s) in DEK responsible for this marked functional impact of DEK in ribosome function, we monitored cellular protein translation in DEK KO cells after re-expression of N-terminal (DEK 1-187, **Figure 3B lane 8**) or C-terminal (DEK 187-375, **Figure 3B lane 10**) fragments or of RBN#3c (DEK 1-375 RBN#3c, **Figure 3B lane 4**) using SUnSET. Whereas DEK1-187 lacked any rescue activity, the C-terminal portion of DEK, containing the two RNA-interacting regions, was sufficient to restore global translational rates. Surprisingly, the RBN#3c mutant expressed as a full-length fusion failed to do so. These results establish the requirement for an RNA binding-competent DEK 178-270 domain, yet not any other DEK region, for unperturbed cellular ribosome function (**Figure 3B, Supplementary Figure 9**).

In line with the marked reduction of translational activity, ribosomal profiling via density ultracentrifugation of cytoplasmic extracts derived from DEK KD or control cells revealed substantial qualitative changes to the cellular ribosomal profile associated with loss of DEK. Whereas relative abundance of 40S small ribosomal subunits (SSU) remained virtually stable (**Figure 3C, lanes 3-5**), substantial reduction of polysomes (**Figure 3C, lanes 10-16**), 80S ribosomes (**Figure 3C, lanes 7-9**), and of 60S large ribosomal subunits (LSU) was seen in DEK KD cells (**Figure 3C, Supplementary Figure 10**) and HeLa DEK KO cells (**Supplementary Figure 11**). Impact of DEK on the large ribosomal subunit was further supported by monitoring RPL19, a prominent component of the LSU, via immunofluorescence and Pre-ribosome Sequential Extraction (PSE). DEK KO or KD cells showed increased nucleolar accumulation and delayed extractability of RPL19 in a DEK-dependent fashion (**Supplementary Figure 12**). Collectively, these findings not only further confirm DEK as a previously unrecognized regulator of ribosome biogenesis, they also specifically outline potential mechanistic involvement of DEK in pathways affecting biogenesis of the LSU, which we investigated next.

Given that the RBN#3c mutant failed to restore translational activity (**Figure 3B**) along with markedly reduced total amounts of CLIP’ed ribosomal RNA (**Figure 2B**), we hypothesized that interaction of DEK with rRNA may represent a dominant mechanistic contribution of DEK in the context of cellular ribosome biogenesis.

### DEK interacts with pol-I transcribed rRNA and broadly affects transcription and processing thereof

To test this hypothesis, we re-analyzed our CLIP data which indicated preferential binding of DEK to 28S rRNA (**Figure 4A-C**). This was further substantiated by RIP-qPCR identifying enrichment of 47S and 28S rRNA species in DEK wt, yet to a much lesser extent in the RBN#3c samples (**Figure 4D**). Moreover, detailed profiling of rRNA reads revealed specific accumulation at the 5’ end of 28S rRNA in DEK wt (**Figure 4E**). This indicates direct interaction of DEK with rRNA and suggests up or downstream roles in the rRNA processing pathway, which may entail effects on transcriptional regulation and/or processing.

**Figure 4.**
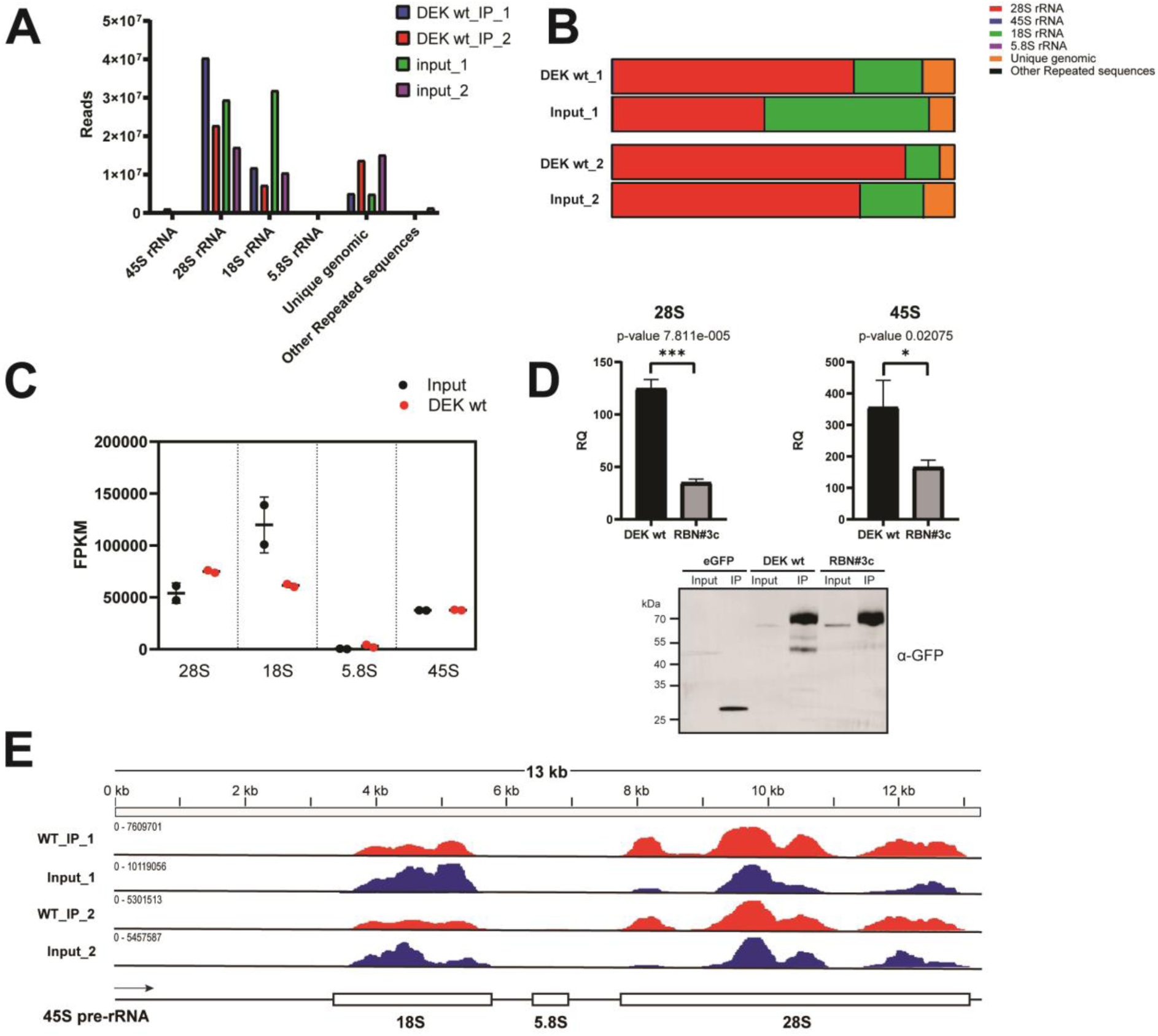
DEK associates with pol-I transcribed rRNA. A. Histogram showing read counts for each rRNA biotype cross-linked to DEK wt, with two biological replicates, DEK wt_IP_1 in blue and DEK wt_IP_2 in red, compared to corresponding inputs (input_1 in green and input_2 in purple); **B.** Stacked bar charts indicate the fraction of each rRNA subtype, the reads uniquely mapped to the human genome and other repeated sequences; **C.** Taking sequencing depth and gene length into consideration, Fragments Per Kilobase Million (FPKM) of each rRNA subtype derived from two DEK wt CLIP samples (in red) and the corresponding inputs (in black) was analyzed; **D.** RNA transcripts immunoprecipitated with eGFP, eGFP-DEK wt or eGFP-RBN#3c, along with their corresponding input RNAs were purified, reverse transcribed and quantified by qPCR using primers annealing to 28S (top left) and 45S rRNA (top right). An aliquot of each sample was subjected to immunoblotting with GFP-specific antibodies to assess expression and immunoprecipitation efficiency of DEK wt and RBN#3c (bottom). Note one fourth of eGFP input sample was loaded on SDS-PAGE to avoid overexposure. Data from three biological replicates were used to calculate p values. **E.** DEK CLIP reads (in red), and input reads (in blue) mapped to human 47S pre-rRNA show a specific binding peak of DEK wt to 5’end of 28S rRNA.

Conversion of precursor rRNA into mature forms destined to be assembled into the two ribosomal subunits involves an extensive sequence of processing steps with the assistance of hundreds of processing factors in mammals. Although numerous variations in timing and processing among species exist, we focused on the predominant 47 S rRNA precursor processing pathway in HeLa cells (**Figure 5A**, adapted from ^54^). Typically, maturation of this polycistronic precursor rRNA starts co-transcriptionally with generation of 18S rRNA by exonucleolytic and endonucleolytic cleavages at 5’ETS and ITS1 followed by trimming of flanking sequences of 5.8S, and removal of the ITS2 spacer sequence is initiated to allow for maturation of 28S and 5.8S rRNA ^72^. Probes, specifically annealing to certain intermediate processing products, allowing for systematic assessment of potential changes in this pathway are schematically listed in **Supplementary Figure 13**.

**Figure 5.**
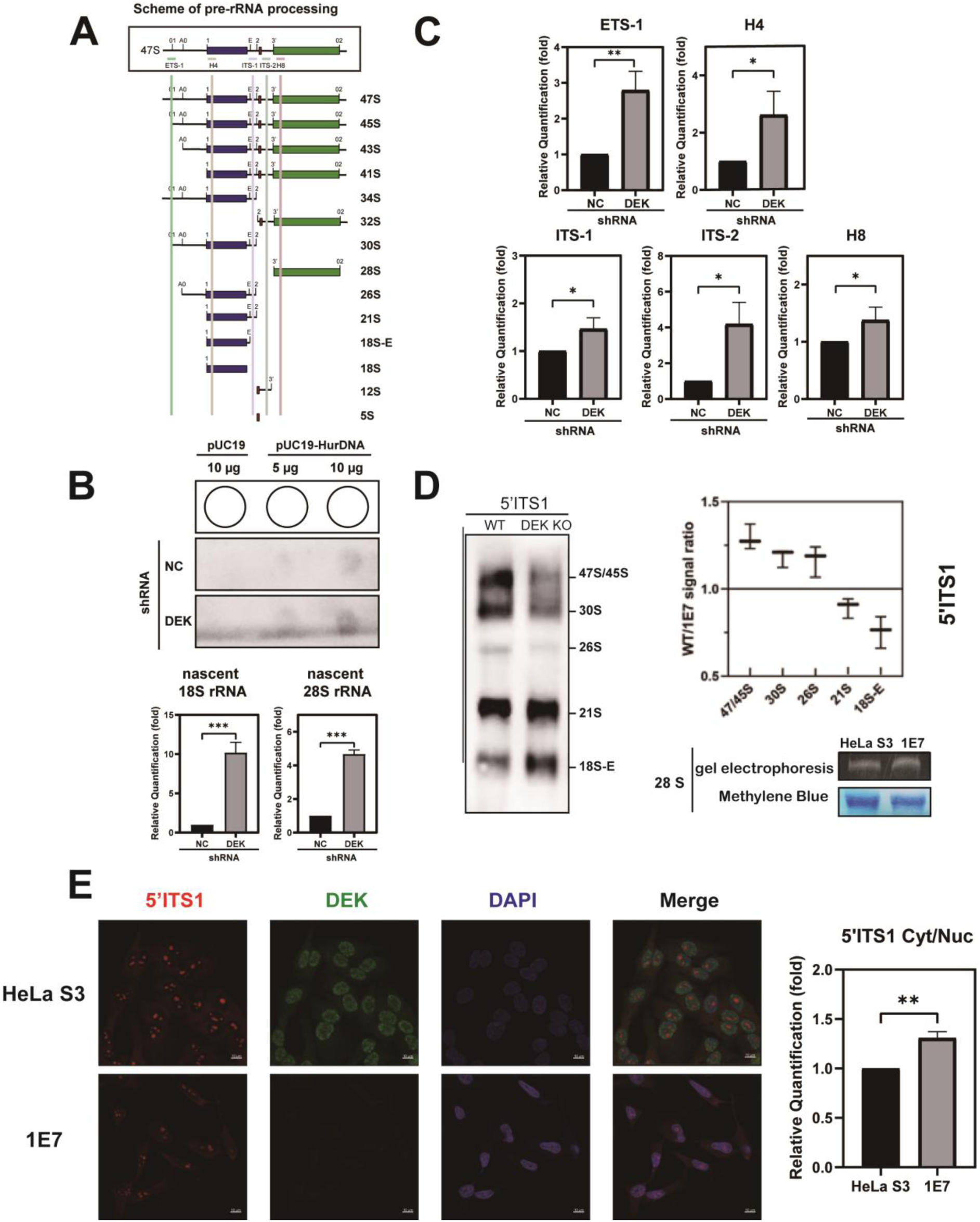
DEK depletion results in markedly increased transcription of pol-I-transcribed pre-rRNA and deficient processing thereof. A. Schematic overview of the primary 47S pre-rRNA processing pathway in HeLa S3 cell (adopted from ^54^). The pre-rRNA is transcribed by RNA polymerase I as a single 47S polycistronic RNA precursor. A series of cleaving steps are required to obtain mature individual rRNA species. **B.** Nuclear run-on assay. Lysates derived from stable HeLa S3 control and DEK knockdown cells were chilled to cause polymerase pausing and restarted with the presence of biotinylated uridine 5’-triphosphate. The labelled nascent RNA products were hybridized with the plasmid containing 47S pre-rRNA gene (pUC19-HurDNA) and pUC19 as a control. An aliquot of labelled transcripts was subjected to immunoprecipitation using Streptavidin magnetic beads followed by qPCR with primers designed for 18S, 28S rRNA gene and Actin for normalization. Delta Ct (CT_18S/28S_ – CT_Actin_) was calculated among three biological replicates (n=3). Student’s t-test (Two-tailed) was applied to the result, ***p < 0.001. **C.** Quantitative analysis of various steady-state rRNA species by RT-qPCR in stable HeLa S3 control and DEK knockdown cells using the primers indicated in (**A**). Graphs represent data from three biological replicates (n=3). Student’s t-test (Two-tailed) was applied to the result, *p < 0.01; **D.** Northern blot analysis. Total RNA from HeLa S3 wt and DEK KO cells was extracted, separated by 1% formaldehyde-agarose gel and transferred to a nylon membrane. After hybridization with biotinylated 5’ITS1 probe, the targeted RNA species with various sizes can be detected by chemiluminescence (left panel). Shown is one representative blot from three independent experiments. For quantification, signals from the three blots were converted to binary mode, followed by measurement of each band by ImageJ. The average signal ratio of each band (n=3) was calculated (right upper panel). To ensure the same amounts of total rRNA from indicated cell lines were loaded on the gel and equally transferred to the membrane, 28S rRNA was examined and detected by GelRed staining of the agarose gel and Methylene Blue staining of the membrane (right lower panel); **E.** HeLa S3 wt and 1E7 cells were hybridized with Cy5 labelled probe annealing to pre-rRNA transcripts containing the sequence complementary to 5’ITS1 (red), followed by immunostaining with monoclonal antibodies specific to DEK (green). DNA was stained with DAPI (blue). Images were taken by a Zeiss LSM880 confocal microscope. Scale bar: 10 μm; For quantitative analysis, the DAPI channel was recognized by the macro of ImageJ as the nuclei. The threshold-mask was then adjusted to 40 to separate signals of the cytoplasmic 18S/28S rRNA from background. Average pixel intensity and pixel coverage were measured giving rise to relative intensities from cytoplasm and nucleus. The ratios of cytoplasmic to nucleic intensities were calculated with approx. 40 cells from four images taken for each measurement. Student’s t-test (Two-tailed) was applied to the result, **p < 0.01.

Interestingly, as DEK depletion resulted in broadly altered nucleolar morphology (**Supplementary Figure 14**), indicative of general nucleolar/ribosomal stress, we first examined the impact of DEK on rDNA transcription by monitoring the abundance of nascent rRNA transcripts in nuclear run-on assays (**Figure 5B**). Equal numbers of nuclei from DEK KD or control cells were isolated, chilled, and polymerase stalling was lifted in the presence of biotinylated uridine 5’-triphosphate (UTP), followed by isolation of total nuclear RNA. Hybridization of resulting RNA to the 5’ end of the 47S rRNA gene (pUC19-HurDNA), or an empty plasmid (pUC19 empty vector) spotted on a membrane, followed by detection via streptavidin antibodies, indicated a relatively higher transcription rate of 47S precursor rRNAs in the absence of DEK. To quantify this effect, biotinylated RNA from nuclear run-on assays was additionally captured by magnetic streptavidin-coated beads, purified, and subjected to RT-qPCR. Consistent with the result from the in-situ hybridization, DEK KD cells displayed a ∼ 4-fold higher transcriptional rate as compared to the control. Moreover, fluorescence *in situ* hybridization (FISH) using Cy5 labelled probes showed uniformly increased signals derived from mature18S and 28S in cytosol or nucleus in DEK KD cells (**Supplementary Figure 15**), in further support of increased transcriptional activity upon DEK depletion.

As rRNA transcription and processing are intimately intertwined, we next assessed major steps along this dynamically regulated processing pathway by qPCR using a set of primers pairs specifically probing across defined sections of the 47S precursor rRNA transcript (**Figure 5C**). The “ETS-1” primer pair detects the first cleavage site of 5’ external transcribed spacer (5’ETS) for estimation of the half-life of 47S pre-rRNA, “H4” anneals to 18S containing rRNA intermediates, “ITS-1” detects the internal space between 18S and 5.8S and “ITS-2” senses the second internal spacer between 5.8S and H8 in rRNA species containing 28S rRNA. To minimize fluctuations due to varying knockdown efficiencies after siRNA transfection, qPCR was performed with RNA extracted from stable cells expressing shRNAs targeting *DEK* ^8^. As expected, knockdown of DEK resulted in significantly higher expression of corresponding rRNA species across all five primer pairs (**Figure 5C**), which was also visible, albeit to varying degrees, in a melanoma cell line (UACC-257) and in primary fibroblasts (**Supplementary Figure 16**), suggesting a universal rather than cell-type specific effect of DEK on rRNA abundance.

To further detail changes in rRNA processing steps, total RNA extracted from HeLa S3 wt and DEK KD was subjected to northern blot analysis with probes complementary to 5’ITS1 (**Figure 5D**). Similar to results obtained from HeLa cells treated with siRNAs ^54^, quantification revealed reduced presence of rRNA species at early stages (47/45S, 30S and 26S) yet accumulation of 21S and 18S-E rRNA intermediates when DEK was depleted. To further examine this observation, we performed FISH using the 5’ITS1 probe labelled with Cy5 at 3’end (**Figure 5E**). As expected, the wt control cells exhibited accumulation of fluorescence signals in the nucleolus, indicating rRNA intermediates at early stages before export in the form of 18S-E rRNA, whereas only a faint signal can be detected in cytoplasm. Upon depletion of DEK, a fraction of the signal shifted from the nucleolus to cytoplasm, which is in line with the blotting data showing accelerated production of 18S rRNA flanked with a fragment of ITS1 in the cytoplasm.

Taken together, these data indicate a pleiotropic role for DEK in transcriptional regulation of rDNA transcription and in facilitating late stages of pol I-transcribed rRNA transcripts encompassing 18S, 5.8S and 28S rRNAs. Although these rRNAs account for the majority of rRNA, 5S rRNA synthesized by pol III from other chromosomal loci in the proximity of the nucleolus ^73^ plays essential roles in facilitating communication among regions of the large ribosome ^74, 75^. To obtain a comprehensive picture of DEK-rRNA interactions we thus next examined a potential connection of DEK to 5S rRNA.

### Direct interaction of DEK with 5S rRNA is involved in 5S RNP formation and response to ribosomal stress via the 5S RNP-MDM2-p53 axis

Mapping of DEK wt and RBN#3c CLIP reads to the 122 bp *5S rRNA* gene revealed a higher density of binding tags in DEK wt versus RBN#3c (**Figure 6A**). To investigate if DEK directly interacts with 5S rRNA, we applied CLIP followed by a tandem purification procedure using HEK293 cells expressing a dual affinity tagged DEK fusion (DEK-His-GFP) or His-GFP as control. After sequential affinity purification via GFP-trap and Ni-NTA agarose, captured rRNAs were subjected to analysis via northern blot using 5.8S and 5S rRNA-specific probes. While the control returned only faint signals, DEK-His-GFP efficiently precipitated 5S rRNA, supporting direct interaction between DEK and this component of the large ribosomal subunit (**Figure 6B**).

**Figure 6.**
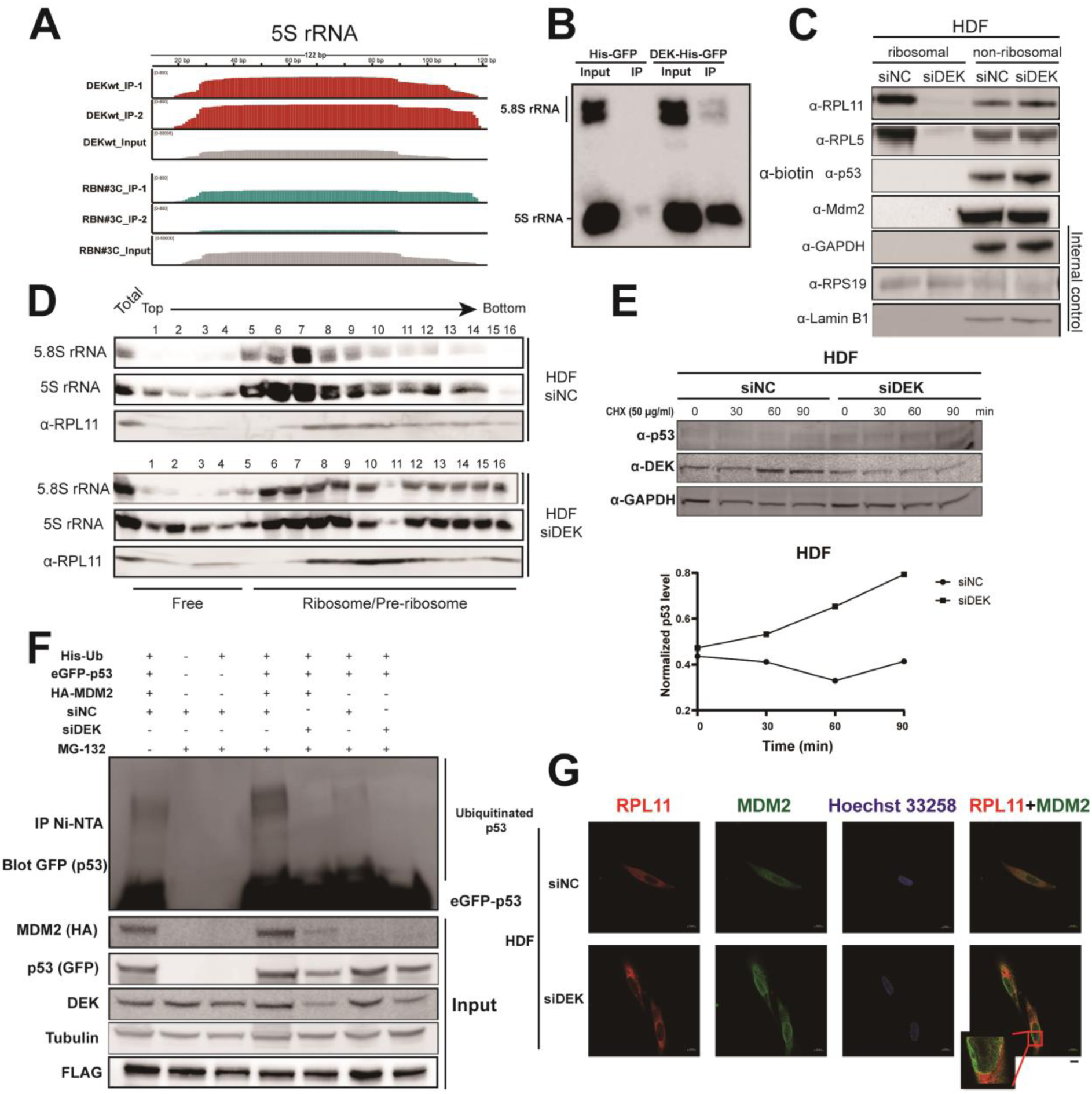
DEK interacts with 5S rRNA and regulates stress response via the p53-MDM2 pathway. A. CLIP reads from DEK wt (in red) mapped to human 5S pre-rRNA show extensive enrichment when compared to RBN#3c (in cyan) and input reads (in gray). **B.** Two-step immunoprecipitation was carried out in HEK293 cells expressing a dual affinity tagged DEK fusion (DEK-His-GFP) or His-GFP as a negative control. The captured RNAs along with inputs were isolated and separated by denaturing polyacrylamide gels followed by Northern Blot with probes annealing to 5S rRNA, and 5.8S rRNA as an internal control. **C.** Primary human dermal fibroblasts (HDF) cells transfected with siRNA targeting *DEK* or control siRNA for 72 hours were harvested, and ribosomal and non-ribosomal fractions were enriched by ultracentrifugation. Resulting samples were analyzed by immunoblotting with the indicated antibodies. GAPDH, RPS19 and Lamin B1 served as internal controls; **D**. Extracts from HDF cells as described in (**C**) were analyzed by sucrose gradient ultracentrifugation followed by detection of 5.8 S or 5 S rRNA by Northern blotting or RPL11 by immunoblotting. **E**. Time-lapse examination of p53 degradation dependent on DEK. HDF primary cells transfected with siRNA targeting DEK or a negative control siRNA for 72 hours were seeded one day before treatment. Cells were treated with 50 µg/mL cycloheximide (CHX) and collected at the indicated time points and analyzed by immunoblotting with the indicated antibodies. The signal intensities were measured by ImageJ and plotted as the intensities of p53 normalized to GAPDH (Supplementary Figure 20). **F.** *In cellula* ubiquitination assay. HEK293 cells transfected with the indicated plasmids were treated with MG132 to a final concentration of 20 µM followed with immunoprecipitation with Ni-NTA beads. IP samples were analyzed by immunoblotting with antibodies specific to GFP and inputs were analyzed with HA-, GFP-, and DEK-specific antibodies. Tubulin was used as an internal control and FLAG as a transfection control; **G.** Immunofluorescence of HDF cells transfected with siRNA targeting DEK or a negative control siRNA. Endogenous RPL11 (red) and MDM2 (green) were detected by confocal microscopy using a Zeiss LSM880 microscope. Scale bar: 10 µm.

Interestingly, 5S rRNA is not only an integral part of the complete ribosome, yet also forms a stable assembly intermediate in concert with RPL11 and RPL5, the so called 5S RNP. Impairment of ribosome biogenesis can cause redirection of un-incorporated 5S pre-ribosomal complexes. This results in non-ribosomal 5S RNP binding to MDM2 which blocks its ubiquitin ligase function and thus resulting in p53 stabilization and accumulation - a pathway recently termed as the impaired ribosome biogenesis checkpoint (IRBC) ^76^. As downregulation of DEK itself resulted in a ribosomal stress phenotype (**Supplementary Figure 14)**, we first monitored if tampering with cellular DEK levels may influence factors involved in the IRBC pathway. Indeed, altering cellular DEK expression in primary fibroblasts resulted in mutual reciprocal changes to p53, p21, RPL11 and RPL5 protein levels (**Supplementary Figure 17A**), and DEK knockdown in UACC-257 melanoma cells, resulted, as expected ^8^, in cellular senescence along with upregulation of p53 and RPL11 (**Supplementary Figure 17B**). These findings are reminiscent of an earlier body of work ^8, 12, 16^, collectively highlighting p53-dependent, yet also independent, roles for DEK in cellular proliferation, apoptosis, and senescence. To investigate if DEK transmits such functions, at least in part, via 5S RNP and the IRBC pathway, we separated bulk ribosomal and non-ribosomal fractions derived from cell lysates of control or DEK KD cells on sucrose gradients via ultracentrifugation. This showed marked and exclusive increase of RPL5 and RPL11 in ribosomal-free fractions in cells with reduced expression of DEK (**Figure 6C and Supplementary Figure 18)**. A more detailed fractionation approach further confirmed increased presence of free RPL11 and 5S rRNA in the absence of DEK (**Figure 6D**), an effect reminiscent of 5-FU treatment of cells (**Supplementary Figure 19**). Together, these results establish DEK as regulator of 5S RNP. As loss of DEK triggers ribosomal stress and may thus activate the IRBC via the 5S RNP-MDM2 axis we investigated such a possibility next.

Cycloheximide time course assays in primary fibroblasts showed that an increase in p53 protein levels is due to the longer half-life at post-translational level in DEK-depleted cells (**Figure 6E, Supplementary Figure 20**). Although similar phenomena have been observed in HeLa cells and concluded to be the consequence of the regulation of DEK on HPV E6 present in the cells ^16^, we next examined the possibility of MDM2, an E3 ubiquitin ligase, -mediated ubiquitination on p53 stabilization upon DEK depletion. In *in cellula* ubiquitination assays, co-overexpression of His-Ubiquitin, HA-MDM2 and eGFP-p53 in HEK293 produced intense ladders representing ubiquitinated p53 products after MG132 treatment as expected (**Figure 6F**). However, cells transfected with the same plasmids but with siRNA mediated DEK KD showed lower intensity of ladders, indicating decreased ubiquitination of p53. Meanwhile, down-regulated MDM2 expression was observed upon depletion of DEK under the condition of the same transfection efficiency (see FLAG expression as a control). These results demonstrated that depletion of DEK appeared to inhibit MDM2-mediated ubiquitination of p53, thus preventing p53 from degradation.

Having confirmed the accumulation of each component of 5S RNP in ribosomal-free pools upon DEK-depletion induced ribosomal stress, we then questioned the direct association of 5S RNP with MDM2, which was proposed to hinder its function on the ubiquitination of p53. Immunofluorescence staining revealed the translocation of a noticeable portion of RPL11 and MDM2 from the cytoplasm to nucleoplasm and the resulting colocalization of these two endogenous proteins on the border of nucleoplasm can be observed in DEK depleted HDF cells, supporting the notion that the MDM2-mediated p53 activation in DEK-depleted cells is probably through the IRBC pathway (**Figure 6G**).

## Discussion

Herein we provide evidence substantially expanding knowledge about the nucleic acid interaction capabilities of the DEK oncogene. Applying crosslinking and RNA immunoprecipitation followed by next-generation sequencing (CLIP-seq) and using a newly established DEK mutant attenuated in its ability to bind to RNA, we now not only further confirm DEK as a mRNA-binding protein at the cellular level, but also reveal a connection between DEK and ribosome biogenesis via interactions with 28S and 5S rRNA. Consequently, DEK depletion resulted in substantially reduced translational activity and qualitative changes to cytoplasmic ribosomes, particularly to the 60S large ribosomal subunit, which can be exclusively rescued by overexpression of C-terminal, rRNA-binding competent DEK. Mechanistically, we provide evidence of broad involvement of DEK in aspects of pol I-mediated rRNA transcription, processing, and localization, and, of particular note, identified specific association of DEK with pol-III transcribed 5S rRNA. Via this newly identified association with the 5S RNP complex, we report functional involvement of DEK in the Impaired Ribosome Biogenesis Checkpoint (IRBC).

An array of *in vitro* and *in vivo* studies over the past three decades highlighted functions of DEK in aspects of chromatin biology. Yet, quickly after its initial cloning an involvement of DEK in cellular RNA biology became evident ^19, 22, 30^. Even though early studies, reporting DEK as a component of exon junction complex (EJC) ^39–41^, were initially challenged ^42–44^, it is now clear that DEK is involved in a number of aspects of RNA metabolism ^77^, with mRNA proofreading, roles in intron retention ^78, 79^ and alternative splicing ^39, 80^ being prominent examples. Despite these studied mRNA-related functions, molecular details of how DEK interacts with RNA in cells remain under-studied, in part attributable to lack of any conventional and readily identifiable RNA binding domain(s) in DEK.

Yet, based on the unbiased RNA-interactome data sets and DEK-centric studies mentioned above, our own previous research and the CLIP data presented here, it is evident that DEK interacts with coding mRNA, rRNA and other RNA biotypes, such as lncRNA or snoRNA. Such DEK-RNA interactions are mediated predominantly via the C-terminal portion of DEK (187-375). This C-terminal half encompasses a lysine and arginine-rich intrinsically disordered region (IDR) (187-250) and the already well-described C-terminal DNA binding and multimerization domain (270-375), containing a winged helix-like structure reminiscent of transcription factors ^66^. Interestingly, disordered protein sequences often function as non-canonical RNA binding domains in a high percentage of human RBPs as recently revealed by many RNA-omics studies ^47 48^. The presence of two RNA binding domains in DEK expands the complexity of how DEK and RNA/DNA molecules may interact, including possibilities of regulation through post-translational modifications as most phosphorylation sites on DEK are clustered in this region. Binding of RNA molecules to DEK may cause co-folding of these two regions, which then may function as an integrated particle. Alternatively, it is also possible that these two regions manipulate different subsets of RNAs individually. Yet, these questions warrant further detailed analysis beyond the scope of this work.

CLIP-seq data from wt full length DEK revealed 266 common binding sites on coding RNAs mapped to 126 unique genes predominantly belonging to GO pathways relating to translational regulation and ribosome biology. DEK binding to these transcripts occurs preferentially at the 3’UTR and on purine-rich sequences, thus in support of functions for DEK in pre-mRNA splicing. In contrast, the mutant RBN#3c showed preferential binding to consecutive adenine sequences, indicating potential association with poly(A)-tailed mRNA transcripts. Taken together, these data may indicate differential binding patterns of these two RNA binding regions, with the major function of DEK 187-270 modulating ribosome biogenesis-related pathways via splicing related events on coding mRNA. Indeed, many CLIPed mRNAs from DEK wt samples, coding predominantly for proteins relevant to ribosome biology, were recently identified with altered splicing patterns upon DEK depletion ^80^. DEK 270-375, in contrast may potentially regulate mRNA stability, export and/or translational initiation via its poly(A)-binding domain. Additionally, except for the discovery of mRNA and rRNA, other RNA biotypes, such as lncRNA or snoRNA, also hint at versatile functions of DEK across cellular pathways. Six lncRNA genes with enriched binding sites among two biological CLIP-seq DEK wt replicates have been found (**Supplementary Table 3**), with CCAT1 and NORAD being well-studied amongst them. CCAT1, overexpressed in a variety of malignancies, associates with CTCF and modulates, amongst others, chromatin loop formation between the promoter and enhancer of c-*Myc*, thus lifting its expression ^81, 82^. NORAD, activated by DNA damage and also identified as an oncogene ^83^ has roles in mRNA degradation of PUMILIO-specific mRNA encoding proteins related to genome stability ^84^. Moreover, interactome analysis revealed an essential role of NORAD in NARC1 (NORAD-activated ribonucleoprotein complex 1) assembly, which consists of TOP1, ALYREF and PRPF19-CDC5L ^85^. Interestingly, both ALYREF and DEK were identified as components of the exon junction complex (EJC) previously and as joint autoantibodies in systemic lupus erythematosus, suggesting concerted functions ^86, 87^. Although only two common snoRNAs (SNORA10 and SNORD133) have been found cross-linked to DEK among two DEK wt replicates, nine snoRNA species were captured in total (**Supplementary Table 4)**. As snoRNAs have critical roles in rRNA folding and interaction with ribonucleoproteins via guiding modification enzymes to specific nucleotides ^88^, DEK may be involved in aspects of these complex pathways relating to ribosome biogenesis. Together, these new data suggest intricate functions of DEK 187-375 and future studies assessing the dual nucleic acid binding behavior of this region will certainly aid in further dissecting the molecular functions of DEK.

Through comparative bioinformatics analysis of CLIP’ed RNA between DEK wt and RBN#3c, we surprisingly uncovered ribosomal RNAs cross-linked to DEK, with overrepresentation of the 5’end of 28S rRNA transcribed by RNA polymerase I (pol I) and 5S rRNA transcribed by pol III, mediated by the DEK187-250 intrinsically disordered rRNA interaction region. Although connections of DEK to ribosome biology have been uncovered in two DEK-centric interactome studies at the proteomic level ^56, 89, 90^, no clear ribosome-related cellular pathway has been ascribed to DEK thus far. To this end, we first confirmed a critical role for DEK in global biosynthesis of cytoplasmic ribosomes that are responsible for cellular protein translation. Interestingly, rescue experiments highlighted that the C-terminal portion of DEK (DEK187-375), harboring rRNA-interacting activities in DEK187-250, but not DEK1-187, as crucial for this ribosome-related function of DEK (**Figure 4**). Based on the hypothesized function of DEK in promoting rRNA transcription through direct binding to the RNA helicase DDX21 ^56^ and as a potential pre-rRNA processing candidate proposed by a genome-wide screening study ^54^, we then examined the impact of DEK on transcriptional onset and subsequent processing of pre-47S rRNA, respectively (**Figure 5**). Strikingly, both scenarios are considerably altered upon depletion of DEK, while its modulation of rDNA transcriptional rate contributes more substantially to higher expression level of mature pol I-transcribed rRNAs at a steady-state according to the total RNA quantitation evaluation. Considering the role of DEK in (hetero)chromatin maintenance ^30, 38, 91^, we suspect that such higher rDNA transcriptional rates are potentially due to loss of heterochromatic silencing of extra copies of rDNA in the genome, which are essential for recombination repair in response to DNA damage ^92^. Excessive rDNA transcription was reported to be toxic to cells by preventing repair of DNA replication forks by failing to provide templates for homologous recombination frequently occurring to the massive rDNA repeats ^92, 93^. Surprisingly, DEK depletion induced hyper-active rDNA transcription yet did not give rise to excessive ribosome production, as identified by others who demonstrated upregulated pol I transcription rate through oncogenic stress and eliciting an increased protein synthesis rate for sustaining excessive cell proliferation ^60^. On the contrary, the dual roles of DEK in 47S rRNA precursor metabolism results in decreased ribosome abundance with lower translational efficiency, a consequence of defective pre-ribosomal intermediates containing aberrant pre-rRNAs along with deficiencies in transport thereof. Such nucleolar stress typically results in a coordinated stress response via activation of the p53 pathway.

Unlike ribosomopathies, directly caused by mutation of ribosomal proteins (RPs), rRNAs or ribosome biogenesis factors (RBFs), aberrant expression of oncoproteins or deregulation of oncogenic pathways upstream of specific ribosome biogenesis steps and resulting in defective ribosomes is often associated with tumorigenesis and cancer development, which has been extensively reviewed by others ^60, 61, 94^. How does the involvement of DEK in ribosome biogenesis contribute to its pro-oncogenic role through cellular pathways that have been well investigated, such as p53 ^8, 12, 16^, mTOR ^17^ or Vascular Endothelial Growth Factor (VEGF) ^95^? Prompted by its binding to 5S rRNA as revealed through our RNA interactome analysis (**Figure 6**) and the interaction with ribosomal proteins (RPL5 an RPL11) as revealed in the DEK interactome ^56^, we investigated a connection with the IRBC checkpoint as a secondary consequence of impaired ribosome biogenesis which involves p53 stabilization in response to cellular stress through the 5S RNP-MDM2 pathway. A number of studies have reported upregulation of p53 upon DEK depletion in both immortalized cell lines and human primary cells ^8, 12, 16^, collectively pointing to a regulation of p53 by DEK at the post-transcriptional level ^16^. Consistent with this, our study revealed that in E6/7-free cells, DEK-depletion induced accumulation of p53 is a result of blocked MDM2-mediated ubiquitin conjugation, thereby preventing subsequent proteasomal degradation (**Figure 6**). Moreover, our present study provides evidence that the inhibitory effect of MDM2 is exerted by its interaction with 5S RNP in the free form of the (pre-)-ribosomes when DEK is depleted, which supports the notion that 5S RNP is a key mediator of IRBC pathway coupling ribosome biogenesis and p53 activation upon cellular stress. Most notably, activation of the 5S RNP-MDM2-p53 axis upon DEK depletion represents a yet unrecognized pathway of DEK involvement in senescence ^12^. As influence of DEK on all components of the 5S RNP was observed, it can be speculated that DEK may function as an accessory protein in this subcomplex facilitating its localization or incorporation into the large ribosomal subunits under normal conditions. Taken together, two pathways of how DEK is involved in 5S RNP-mediated p53 activation have been proposed here (**Figure 7**). One is via indirect regulation by DEK in the upstream steps to activate ribotoxic insults to elicit the release of 5S RNP sub-complex in a free form from ribosomes and to be redirected to the MDM2 to elicit the IRBC. On the other hand, DEK is a potential direct player in 5S RNP recruitment to the ribosome, with similar functions to PICT1, RRS1 and BXDC1 that have been well documented ^96^. However, to support this notion, further detailed biochemical experiments are required.

**Figure 7.**
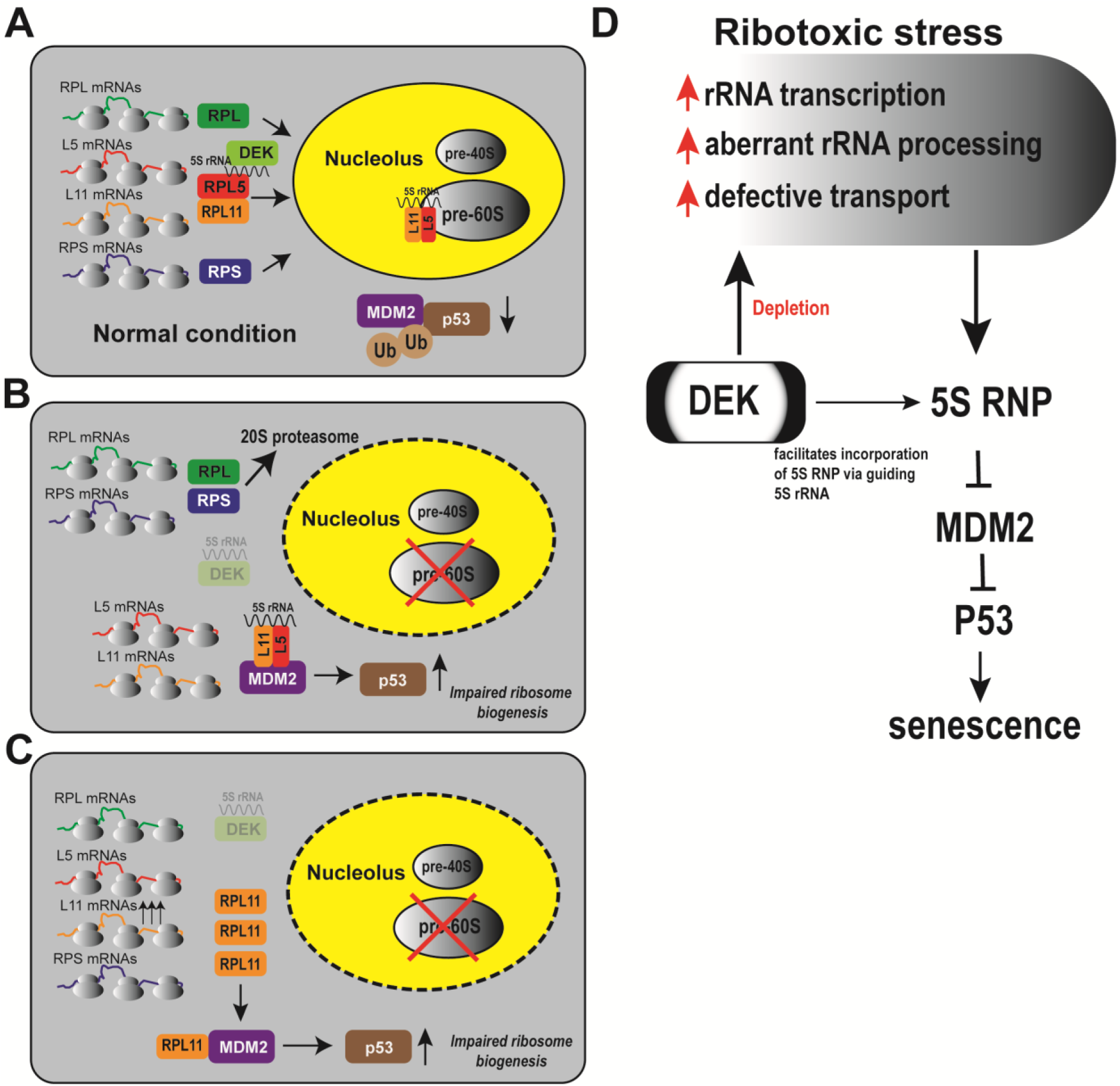
Potential models depicting the roles of DEK in impaired ribosome biogenesis. A. In healthy cells, newly translated ribosomal proteins, including small RPSs for 40S and RPLs for 60S ribosomal subunits, are imported into the nucleolus and subsequently assembled to pre-ribosomal particles together with rRNAs, thereby MDM2 is competent for ubiquitination of p53 leading to its degradation. During the process, DEK exerts the function to facilitate the incorporation of 5S RNP into pre-60S subunits via direct association with 5S rRNA (I); B. Conditions of impaired ribosome biogenesis, leading to disrupted nucleoli and passive release of RPs and ribosomal particles from the nucleolus. Most of the released RPs are degraded by 20S proteasome, while RPL5 and RPL11, two proteins involved in the 5S RNP, are retained through a mutual protection mechanism ^101^. Depletion of DEK results in 5S RNP being freed from the pre-60S subunits, leading to its direct interaction with MDM2 and the blockage of E3 ubiquitin ligase activity towards p53, thus p53 is stabilized (II), **C**. Under certain conditions, RPL11 is probably directly upregulated upon DEK depletion-caused open chromatin at the transcriptional level or indirectly upregulated via its leader sequence (5’TOP) at the translational level, the redundant fraction of RPL11 binds to MDM2 to inhibit its function towards p53 degradation in the absence of nucleolar disaggregation (III). **D.** Simplified graphic summary: The IRBC checkpoint can be elicited by aberrant actions of DEK via two different pathways. One is through DEK depletion-induced ribotoxic stress, including upregulated rRNA transcriptional rate, aberrant rRNA processing with the result of a defective pre-ribosomal particles transport. Consequently, these stressors are thought to activate p53 and drive cells into senescence via 5S RNP (I); Alternatively, DEK serves as an assembly factor of 5S RNP via direct association with 5S rRNA promoting the incorporation of 5S RNP into pre-60S subunits. The loss of DEK may lead 5S RNP to stay in the ribosomal-free pool and interact with MDM2 to inhibit its E3 ubiquitin ligase activity towards p53, leading to p53 stabilization in response to cellular stress (II).

Although it has been generally accepted that the RPL5/RPL11/5S RNA is the predominant sub-complex required for p53 stabilization upon ribosomal stress, questions remain of how 5S RNP senses upstream defects. The earlier assumption is that the nucleolus, as both the primary workshop of ribosome biogenesis and the sensor of ribosomal stress, tends to collapse in response to ribotoxic stress, leading to diffusion of nascent 5S RNP for further incorporation with MDM2 in the nucleoplasm ^62, 97^. In line with this notion, morphologic alternation of nucleoli can be observed in DEK KO cells and enhanced colocalization of endogenous MDM2 and RPL11 in nucleoplasm may imply their direct association for blocking effect on p53 degradation. Other studies also found elevated p53 levels upon impaired ribosome biogenesis under the condition where the nucleolus is undisrupted. The authors claimed that the redundant 5S RNP components targeting to MDM2 are derived from upregulation of mRNAs with a 5’ terminal OligoPrymidine (5’TOP) motif at a translational level ^98, 99^. Interestingly, during the process of DEK knockdown in multiple cell lines, we indeed observed the upregulation of protein expression of RPL11, as one of the top mRNPs, making this explanation also feasible. Thereby, we propose both alternative models explaining the way through which 5S RNP-dependent p53 is elicited upon DEK depletion induced ribosome biogenesis stress (**Figure 7**).

In summary, we have uncovered DEK as a cellular RNA binding protein, with, amongst others, a critical role in regulating pol I-transcribed rDNA transcription and efficient pre-rRNA processing. Furthermore, we linked the molecular function of DEK in ribosome biogenesis to formerly reported roles in p53 activation and inhibition of senescence. Previously, several anticancer compounds have been designed to activate IRBC by inhibiting pol I mediated rDNA transcription, such as CX-5461 and BMH-21 ^60, 61, 100^. Therefore, our observations that DEK is an upstream regulator of rDNA transcription may be instrumental for positioning DEK as a target for therapeutic intervention in cancers. Collectively, our data demonstrate that DEK, a protein best known for its chromatin regulatory function, has previously unrecognized functions in ribosome biogenesis and as a regulator of the RPL5/RPL11/5S rRNA-MDM2-p53 stress response.

## Supplementary Data

Supplementary Data contains 20 Supplementary figures, 3 Supplementary Files and 7 Supplementary Tables.

## Acknowledgements

We thank Joel Mackay (University of Sydney) for pentaprobe plasmids. This work was supported by funds from Xi’an Jiaotong-Liverpool University (RDF-16-02-28 to FK), by the Deutsche Forschungsgemeinschaft (DFG KA 2799/1 to FK), by the START Program of the Faculty of Medicine, RWTH Aachen University (to FK) and by a Wang-Chai Biochemistry and a Start-Up grant from Duke Kunshan University. Funding for open access charge: Duke Kunshan University.

## Author contributions

N.X performed most experiments; K.C performed bioinformatics analyses; M.P established DEK KO cells, cloned LAP-Tag DEK and performed ^35^S incorporation and ribosome profiling experiments; P.L purified GST-DEK fusion from *E. Coli* and performed EMSA; S.Y. and S.G. helped with northern blotting; YL, SD and SI helped with data analysis. N.X and F.K conceived the study and wrote the manuscript with input from all authors.

## Material and Methods

### Cell culture

HeLa S3 wildtype or a TALEN-mediated DEK knock-out cell line (“1E7”,described in ^27, 64^) and UACC-257 melanoma cells ^8^ were cultured in DMEM medium supplemented with 10% FBS and 1% Pen-Strep in a cell culture incubator setting at 37°C, 95% humidity and 5% CO_2_. Cells were split every two or three days by Trypsin-EDTA after washing with PBS. For re-expression of eGFP or eGFP-tagged DEK fusions in 1E7 or other cell lines, an inducible lentiviral system based on the pTRIPZ vector (Open Biosystems) was used as previously described ^27, 64^. Resulting cell lines were selected with 1 µg/ml puromycin (InvivoGen) and maintained under standard conditions (37°C, 5% CO_2_, DMEM + 10% FBS + 1 x Pen/Strep). Expression of eGFP-fusions was induced by addition of doxycycline (1 μg/mL, Thermo) as indicated.

HEK293 Flp-In T-Rex cells with inducible expression of localization and affinity purification (LAP) tagged DEK fusions were created via lentiviral transduction to a single locus as described ^102^. The established cell lines were kept in DMEM medium with 20 µg/mL blasticidine (InvivoGen) and 150 µg/mL hygromycine (BioFroxx). Human Dermal Fibroblast (HDF) primary cells (Procell, China) were cultured in complete medium as supplied by Procell (China) and were split every three to five days when confluency reached ∼70 %.

### Downregulation of endogenous DEK by siRNA and shRNA

siRNA oligonucleotides targeting *DEK* (5’-GAAGGCTAAGCGAACCAA A-3’) at a final concentration of 50 nM were transfected into cells using Lipofectamine 3000 (Invitrogen) according to the manufacturer’s instructions. Cells were collected 72 hours post transfection and the resulting knockdown efficiency was evaluated by RT-qPCR and immunoblotting. For stable knockdown of *DEK* via shRNA, the pLKO.1 vector system delivering the shRNAs was used as previously described ^8^. A 4-day protocol established by the RNAi consortium (https://www.addgene.org/protocols/plko/#E) was used.

### Bacterial Growth Inhibition Screen (BGIS)

Selection of RNA-binding deficient DEK mutants was carried out via BGIS as described in ^27, 64^. Briefly, *E. coli* BL21 (DE3) transformants containing glutathione-S-transferase (GST) tagged protein fusions with mutations generated by error-prone PCR were first plated on LB-ampicillin (AMP, 100 ng/mL) plates. After replica-stamping of colonies to fresh plates containing 0.1 mM IPTG for 3-4 consecutive days, “escaper” colonies were considered as candidates expressing DEK fusions deficient in RNA-binding. After small scale expression and analysis via immunoblot, plasmids from colonies expressing products of proper molecular weight were purified and analyzed by sequencing. RNA-binding activity of resulting mutants was determined *in vitro* by RNA-EMSA with RNA pentaprobes as described previously ^27, 64, 103^.

### CLIP-seq of DEK from HeLa S3 cells

Approximately 5 x 10^7^ HeLa S3 were cultivated in normal medium supplemented with 1 µg/mL doxycycline to induce expression of eGFP-tagged protein fusions for 24 hours. On the next day, cells were washed with pre-chilled PBS and irradiated on ice with 150 mJ/cm^2^ at 254 nm to crosslink RNA and proteins in proximity, followed by scraping and centrifugation at 1,000 x g for 2 minutes at 4°C. The pellet was then resuspended in 1 mL modified iCLIP lysis buffer (50 mM Tris-HCl, pH 7.4, 1% Igepal CA-630, 0.5% SDS, 0.5% sodium deoxycholate) supplemented with cOmplete protease inhibitor cocktail (Roche) and 40 units of RNase Inhibitor (Beyotime) and was kept on ice for 15 minutes. The lysates were passed through a 10 ml syringe with a 30 Gauge needle 15 times to shear genomic DNA. A DNA digestion step followed by addition of 2 units of DNase I (Beyotime) in parallel with 0.1 units of RNase I (Beyotime). The reaction was incubated at room temperature for 15 minutes and put back immediately on ice for 5 minutes. The lysates were then centrifuged at 22,000 x g for 10 minutes at 4°C. The cleared lysates were transferred to a new tube and sonicated with four pulses at 50 % amplitude on ice to decrease the viscosity of lysates, followed by another centrifugation step as before. Two percent of the lysate was removed as input and kept at -80°C until further analysis. For immunoprecipitation, the cell lysates were incubated with 25 μL pre-washed GFP-trap agarose beads (Chromotek) in a cold room for two hours. Beads were then washed once with lysis buffer, twice with high salt buffer (50 mM Tris-HCl, pH 7.4, 1 M NaCl, 1 mM EDTA, 1% NP-40, 0.25% SDS, 0.5% sodium deoxycholate) and one more wash with lysis buffer. After these washing steps, beads were resuspended in 50 μL dephosphorylation buffer (50 mM Tris-HCl, pH 7.9, 100 mM NaCl, 10 mM MgCl_2_) supplemented with 0.5 U/μL of Quick Calf Intestinal Phosphatase (Quick CIP, NEB) to remove 5’ or 3’ phosphates caused by RNase I cleavage. The suspension was incubated at 37°C for 10 minutes and the reaction was stopped by two washes using 1 mL phosphatase washing buffer (50 mM Tris-HCl, pH 7.5, 20 mM EGTA, 0.5% NP-40) and two washes with T4 polynucleotide kinase (PNK) buffer (50 mM Tris-HCl, pH 7.5, 50 mM NaCl, 10 mM MgCl_2_) without DTT. The beads were then resuspended in a 30 μL RNA 3’ End Biotinylation system with 15% PEG, 1 x RNA Ligase Reaction buffer, 40 U RNase inhibitor, 40 U T4 RNA Ligase and Biotinylated Cytidine (Bis)phosphate supplied by RNA 3’ End Biotinylation Kit (Pierce). After incubation at 16°C overnight, the beads were washed three times with washing buffer (50 mM Tris-HCl, pH 7.4, 1 M NaCl, 1 mM EDTA, 1% NP-40, 0.25% SDS, 0.5% sodium deoxycholate) and then resuspended in 70 μL 1 X SDS-PAGE loading buffer and heated at 95°C for 5 minutes to release RNA-protein complexes. The released DEK-RNA complexes were loaded on an 8% HEPES PAGE gel (Yeason) and transferred via a semi-dry blotting system (Biometra) to a 0.45 μm nitrocellulose (NC) membrane with 0.8 mA/cm^2^ for 70 minutes. For RNA isolation, the chosen membrane region was excised with an RNase-free scalpel and immersed in 400 μL PK buffer (100 mM Tris-HCl, pH 7.4, 50 mM NaCl, 10 mM EDTA) supplemented with proteinase K (Beyotime) at a final concentration of 2 mg/mL for 20 minutes at 55°C. Then 400 μL of PK/urea buffer (100 mM Tris-HCl, pH 7.4, 50 mM NaCl, 10 mM EDTA, 7 M urea) was added and incubated for another 20 minutes. RNA was extracted and purified with TRIzol RNA extraction reagent (Thermo) following manufacturer’s instructions. RNA samples were sent to RiboBio (Guangzhou, China) company for cDNA library construction using NEBNext^®^ Ultra™ RNA Library Prep Kit for Illumina (Illumina) and high-throughput sequencing on Illumina sequencing platform choosing PE150 mode to obtain sequence information from both ends.

### Bioinformatics analysis of CLIP-seq data

The raw reads for each CLIP-seq sample were pre-processed by the trimmomatic programme ^104^ applying default parameters for adaptor removal and quality filtering. The filtered reads were then mapped with TopHat (v2.0.13) ^105^ to the human genome (hg19) allowing for 2 mismatches/gaps within the alignment. All uniquely mapped reads were then processed with ANNOVAR ^106^ for functional annotation and Guitar ^107^ was used for genomic distribution analysis. Piranha was used for peak calling. To identify the specific binding sites of DEK wt and RNN#3c, RNA peaks from CLIP and input libraries were firstly generated with Piranha software, then Pyicoenrich was used to detect differential enrichment and to select those RNA peaks in the CLIP library with higher signal when compared to input under the condition of z-scores being higher than 2 ^108^. For annotation and classification of peaks, Homer ^109^ software suite was used with the estimated false discovery rate setting at 0.1%. The enriched motifs within the peak sequences were identified via MEME suite ^110^ with the width between 8 to 12 nt and zero or one occurrence of the motif per sequence (zoops) was set.

### UV crosslinking and two-step immunoprecipitation of 5S rRNA

HEK293 Flp-In T-Rex cells expressing DEK-His-GFP or His-GFP as a negative control were induced using 50 ng/mL doxycycline for 24 hours. Prior to the two-step purification, cells were firstly cross-linked with UV at 150 mJ/cm^2^ and lyzed in 500 μL of RIP-LB buffer (20 mM Tris-HCl, pH 7.5, 150 mM NaCl, 1 mM MgCl_2_, 1 mM CaCl_2_, 0.1% SDS, 1% sodium deoxycholate, 1% Triton X-100) with addition of 20 U RNase Inhibitor (Beyotime), 5 mM PMSF (Beyotime) and 5 mM DTT, and incubated on ice for 10 minutes to complete lysis. Debris was cleared by centrifugation at 16,000 x g for 7 minutes (4°C). The supernatant was kept as cell lysate for further immunoprecipitation. After transferring 100 μL of the cleared lysate to a new 1.5 mL tube as input control, the rest was added to pre-washed GFP-trap suspended in 4.4 mL RIP-DB (10 mM Tris-HCl, pH 7.5, 150 mM NaCl, 0.5 mM EDTA) and incubated with pre-washed GFP-trap agarose beads for 2 hours in a cold room. Beads were then pelleted by centrifugation at 2,000 x g (4°C) followed by five washes with 1 mL of RIP-WB (50 mM Tris-HCl, pH 7.5, 500 mM NaCl, 4 mM MgCl_2_, 0.5% sodium deoxycholate, 0.1% SDS, 2 M urea, 2 mM DTT) and one more wash with 1 mL of cold RIP-LB. The resulting supernatant was incubated with Ni-NTA beads (QIAGEN) under denaturing condition (6M guanidium-HCl) for 3 hours (4°C), followed by three washes with RIP-WB buffer. To release nucleic acids from complexes, beads were suspended in 200 μL of proteinase K buffer (100 mM Tris-HCl, pH 7.4, 50 mM NaCl, 10 mM EDTA) with addition of proteinase K (Beyotime) at a final concentration of 2 μg/μL and incubated at 55°C for 20 minutes. After that, 200 μL of PK/urea buffer (100 mM Tris-HCl, pH 7.4, 50 mM NaCl, 10 mM EDTA, 7 M EDTA) was added and incubated for another 20 minutes at 55°C. The reaction was terminated by addition of Laemmli buffer and boiling at 95°C for 5 minutes followed by Northern blot analysis.

### Surface Sensing of Translation (SUnSET) in human mammalian cells

SUnSET assay was applied to monitor nascent protein synthesis rate ^71^. 0.5 x 10^6^ cells were seeded on 60 mm dishes and grown for 24 hours prior to transfection. On the next day, Lipofectamine 3000 (Invitrogen) was used for transfection of vectors or siRNA following manufacturer’s instructions. 48 hours post transfection, puromycin (10 mg/mL, InvivoGen) was added at a final concentration of 10 µg/mL for 10 minutes. Cells were lysed with 2% SDS followed by immunoblotting using puromycin-specific antibodies (12D10, Sigma).

### 35S-Methionine labeling of cells

Cells were seeded one-day prior labeling in six-well plates. On the following day cells were washed twice with methionine-free medium and then 2 mL methionine-free medium containing dialyzed FCS and 50 μCi 35S-methionine (37 TBq / mmol; 10 mCi / ml) was added to the cells followed by incubation for 2 h at 37°C. Cells were washed twice with ice-cold PBS and 1 mL ice-cold PBS was added and the cells were scraped off the plate. Cells were pelleted at 1500 rpm for 5 min at 4°C. The supernatant was removed, and the cells were resuspended in ice-cold lysis buffer (300 mM NaCl, 1% Triton X-100, 50 mM Tris, pH 7.4, Complete Protease Inhibitor Cocktail, 0.1 mM PMSF) and were incubated for 30 min on ice followed by centrifugation at 4°C. The supernatant was separated via SDS-PAGE, the gel was dried and exposed to an X-ray film. Cells seeded in parallel and not treated with ^35^S-methionine were used to measure total protein concentration.

### Profiling of cytoplasmic ribosomes

4 x 10^6^ cells were seeded two days prior harvest on 150 mm dishes. On the harvest day, collection of cells started after treatment with 100 μg/mL cycloheximide for 10 min in the cell culture incubator. The plates were then placed on ice and washed with ice-cold PBS three times before scraping cells. Cells were counted by a CASY cell counter to ensure the same numbers of cells for each sample were further processed. Cells were collected in 500 μL PBS and mixed with 2.5 mL of ice-cold lysis buffer (0.5% NP-40, 150 mM KCl, 20 mM HEPES, pH 7.6, 5 mM MgCl_2_, 1 mM DTT) containing 100 μg/mL cycloheximide. Cells were kept on ice for 5 minutes followed by centrifugation at 4°C for 10 minutes to separate nuclei and cytoplasm. Equal concentrations of cytoplasm as determined by a BCA kit (Beyotime) were layered on a 15% to 45% (w/v) sucrose gradient supplemented with 100 μg/mL of cycloheximide followed with ultracentrifugation using a swing-out rotor (SW41Ti, Beckman) at 4°C with 40,000 rpm (36,500 x g) for 2 hours. After ultracentrifugation, 800 μL aliquots were carefully taken from the top of the gradient and OD_260_ was measured via Nanodrop for each fraction. This procedure was repeated until the ultracentrifuge tube was empty. For determination of 40S, 60S and 90S ribosomes, RNA was isolated from each fraction with TRIzol (Invitrogen) following manufacturer’s instructions. During the process, 500 pg of External Standard (ExSt or A Luc / CAT) RNA was spiked in each fraction to monitor RNA purification efficiency. RNA was reverse transcribed with Hifair® 1st Strand cDNA Synthesis Kit (YEASEN) following manufacturer’s instructions and PCR was conducted using the cDNA as a template with the primers listed in **Supplementary Tables 5-7**.

### Northern Blot for analysis of rRNA intermediates

For analysis of rRNA processing, the blotting procedure was modified from ^111^. Specifically, total RNA was extracted with TRIzol RNA extraction reagent (Thermo) following manufacturer’s instructions. Three or six µg of purified RNA was separated on a 1.2% agarose-formaldehyde gel after 15 minutes incubation at 70°C for denaturation. RNA was then transferred to pre-wetted Hybond N+ nylon membrane (Beyotime) by capillary blotting system with 10 X SSC buffer (1.5 M NaCl, 0.15 M sodium citrate, pH 7.0) overnight. On the next day, transferred RNA was immobilized by 10-minute baking in an 80°C oven and two times irradiation at 150 mJ/cm^2^ with a UV crosslinker, followed by pre-hybridization in Church buffer (1 % BSA, 1 mM EDTA, 0.25 M phosphatase buffer, pH 7.2, 7 % SDS, 0.9 µg/mL tRNA from brewer’s yeast (Roche)) for 1 hour at 45°C. 3’ biotinylated single-stranded DNA probe (5’ITS1: 5’-cctcgccctccgggctccgttaatgatc-3’) was added for hybridization overnight. The membrane was then washed twice with 0.1 X SSC – 0.1 % SDS [wt/vol] wash buffer (10 minutes each time). Biotin signal was detected following the instruction of Chemiluminescent Biotin-labelled Nucleic Acid Detection Kit (Beyotime). See **Supplementary Tables 5-7** for probe sequences.

### Fluorescence *In Situ* Hybridization (FISH) and Immunofluorescence (IF)

3x10^5^ cells were seeded on 60 mm dishes with a coated glass slide at bottom one day before cell fixation. On the second day, cells were rinsed with PBS three times, then fixation was carried out with 4 % paraformaldehyde (PFA) for 10 minutes at RT. PFA was washed off by one wash with PBS, followed by a 10-minute permeabilization step on ice with 0.5 % Triton X-100 supplemented with 20 mM Ribonucleoside Vanadyl Complexes (RVC) (Beyotime) to protect RNA from degradation. The slide with fixed cells was then transferred to pre-chilled 70 % ethanol for at least 10 minutes until the next step. Hybridization was carried out by treating cells with 5 μg/μL of labelled DNA probe diluted in hybridization buffer (1 X Denhardt’s solution (Solarbio), 10 % BSA, 25 % formamide, 10 mM RVC, 2 X SSC) overnight in cell culture incubator. The next day, hybridization was terminated by one wash with 4 X SSC and two washes with 2 X SSC high salt buffer (5 minutes each time), followed by blocking in 4 % BSA in PBST for 20 minutes in the dark at RT. After removing blocking buffer, primary antibodies diluted in 1 % BSA with the dilution ratio recommended by the manufacturer’s instructions were incubated with cells for 1 hour at RT in the dark. Three washes with PBST followed (3 minutes each time) with a last incubation with secondary antibodies for 1 hour at RT. Cells were rinsed again with PBST and then DNA was stained using either DAPI (Beyotime) or Hoechst 33528 (Beyotime) staining following the manufacturer’s instructions, followed with two rinses in PBST and mounted by mounting medium (Beyotime). Observations were carried out using a Zeiss LSM880 microscope.

### Nuclear run-on assay

1 x 10^7^ – 10^8^ cells were collected in a 15 mL reaction tube. The cell pellet was resuspended in 5 mL of pre-cooled NP-40 lysis buffer (10 mM Tris-HCl, pH 7.9, 10 mM NaCl, 3 mM MgCl_2_, 0.1 % NP-40) followed by incubation on ice for 5 minutes. Nuclear pellet was sedimented by centrifugation at 200 x g for 10 minutes (4°C) and stored in 500 μL of nuclear freezing buffer (10 mM Tris-HCl, pH 7.9, 0.1 mM EDTA, 3 mM MgCl_2_, 40 % glycerol). Polymerase pausing was carried out by freezing at -80°C overnight. On the next day, 225 μL of the mixture was combined with 60 μL of 5 X run-on buffer (25 mM Tris-HCl, pH 8.0, 12.5 mM MgCl_2_, 750 mM KCl) supplemented with 40 U of RNase inhibitor (Beyotime) and 2 μL of biotin RNA labelling mix (Roche) and incubated at 37°C for 15 minutes for the purpose of elongation and biotinylated UTP integration. Genomic DNA was then digested by adding 20 μL of 10 mM CaCl_2_ to a final concentration of 200 μM and 200 U DNase I (Thermo) and incubated at 30°C for 5 minutes to increase its viscosity, followed by incubation with 100 μg of proteinase K (Beyotime) at 42°C for 30 minutes. RNA was extracted by TRIzol and further processed for detection by Northern Blot and RNA pull-down followed by qPCR.

For Northern Blot analysis, approximately 10 μg of DNA plasmid (pUC19 empty vector or pUC19-HurDNA constructed as described by ^112^ linearized by BamHI was precipitated and diluted in TE buffer to a final volume of 400 μL for each dot. They were applied to a positively charged nylon membrane using a dot-blot apparatus and rinsed with 6 X SSC buffer (600 μL) twice. The membrane was then dried in an 80°C oven for 2 hours for fixation. Signals can be detected after hybridization with 20 mL hybridization buffer containing 5 μg of RNA extracted from the previous step.

Newly synthesized RNA was also captured by BeyoMag™ Streptavidin Magnetic Beads (Beyotime) according to the manufacturer’s instructions followed by RNA precipitation, reverse transcription, and RT-qPCR for quantification.

### *In vivo* ubiquitination assay

Cells were transfected with 20 μg of plasmids in total (10 μg His-Ubiquitin, 5 μg eGFP-p53 and 5 μg HA-MDM2). 48 hours post transfection, cells were treated with 25 μM MG-132 for 3 hours before harvest. Cells were then lysed with 500 μL of cell lysis buffer (6 M guanidinium chloride, 0.1 M phosphatase buffer, pH 8.0, 10 mM Tris-HCl, pH 8.0) followed by incubation at room temperature for 30 minutes. 40 μL of the lysate was taken as input control before adding 75 μL of washed Ni-NTA (QIAGEN) beads slurry and incubation for 4 hours in a cold room. After incubation, beads were collected by centrifugation at 2,000 x g for 5 minutes and then washed by buffer (8 M urea, 0.1 M Na_2_HPO_4_/NaH_2_PO_4_, pH 8.0, 10 mM Tris-HCl, pH 8.0), B (8 M urea, 0.1 M Na_2_HPO_4_/NaH_2_PO_4_, 10 mM Tris-HCl, pH 6.3) and C (8 M urea, 0.1 M Na_2_HPO_4_/NaH_2_PO_4_, 10 mM Tris-HCl, pH 8.0, 0.1% Triton X-100) sequentially. The ubiquitinated proteins were finally suspended in 50 μL elution buffer and eluted with 5 minutes incubation with vigorous shaking. Both input and IP samples were subjected to SDS-PAGE and analyzed by immunoblotting with antibodies indicated at the respective figure.

### Isolation of ribosomal and non-ribosomal cytoplasmic and nuclear fractions

Five 10 cm plates of cells with ∼80 confluency were collected and resuspended in 3 mL of low salt buffer (LSB, 10 mM HEPES-NaOH, pH 7.5, 10 mM NaCl, 2 mM MgCl_2_, 1 mM EDTA) and incubated on ice for 10 minutes followed by centrifugation at 1,200 x g for 5 minutes (4°C). The pellets were resuspended in 3 ml LSB buffer supplemented with NP-40 to a final concentration of 0.3 % and sodium deoxycholate to 0.2 % followed by vigorous vortexing for 30 seconds. After centrifugation at 2,800 x g for 5 minutes, the resulting supernatant was stored as the cytoplasmic fraction with a volume of approximately 3.5 mL. The nuclei pellet was then subjected to lysis with 900 μL of high salt buffer (HSB, 10 mM Tris-HCl, pH 7.2, 0.5 M NaCl, 50 mM MgCl_2_, 0.1 mM CaCl_2_) with addition of 20 U RNase inhibitor (Beyotime) and 150 U DNase I (Thermo) and 10 minutes incubation at RT, followed by centrifugation at 12,000 x g for 10 minutes. The pellet was rinsed with 300 μL of ice-cold nuclear extraction buffer (10 mM Tris-HCl, pH 7.2, 10 mM NaCl, 10 mM EDTA) and centrifuged at 12,000 x g for 1 minute. The resulting supernatants from these two centrifugation steps were combined and defined as nucleoplasmic fraction (with approximately 1.2 mL in total). For isolation of ribosomal and non-ribosomal fractions, samples were layered on a 20 % [wt/vol] sucrose cushion with 0.36 mM cycloheximide, respectively, followed by ultracentrifugation at 149,000 x g for 2 hours (4°C). The resulting supernatant was taken as non-ribosomal part of cytoplasmic or nucleoplasmic fraction, while the pellet was considered as ribosomal particles. They were analyzed by immunoblotting following a protein precipitation procedure.

