## Supplemental Information for "DEK-rRNA interactions regulate ribosome biogenesis and stress response"

**Supplementary Figures 1-20**

**Supplementary Tables 1-8**

**Supplementary Files 1-3 (*individual files not included in this pdf file*)**

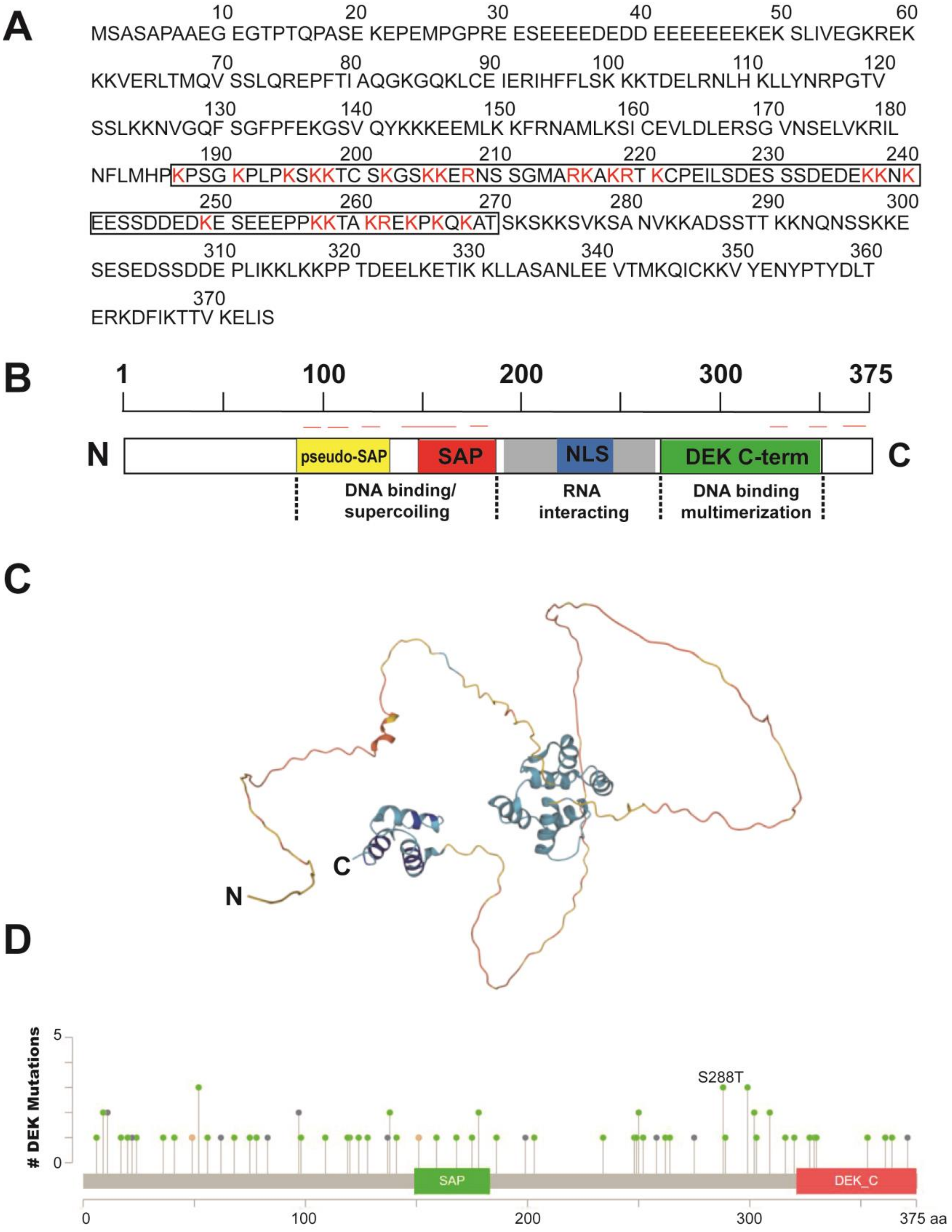


**Supplementary Figure 1. Sequence and structure analysis of the human DEK protein.**

**A.** Amino acid sequence of full length human DEK with the RNA-interaction domain (amino acids 187-270) highlighted by a box. Lysines (K) and arginines (R) are indicated in red. The overall percentage of positively charged amino acids in this region is with 30 % rather high.

**B.** Schematic depiction of human DEK with identified domains indicated in different colours (*yellow:* pseudo-SAP-box; *red:* SAP-box; *blue:* nuclear localization sequence (NLS); *grey*: RNA binding domain *green:* C-terminal DNA-binding domain). The documented functions of each motif or region are shown at the bottom. Red lines on top: positions of α-helices [1, 2].

**C.** Three-dimensional structure of human DEK as predicted by AlphaFold [3]. Accessed June 2024.

**D.** Occurrence of DEK mutations identified in 10,967 samples derived from 32 studies summarised by cBioPortal (https://www.cbioportal.org), indicating no prominent mutational hotspots (accessed June 2024).

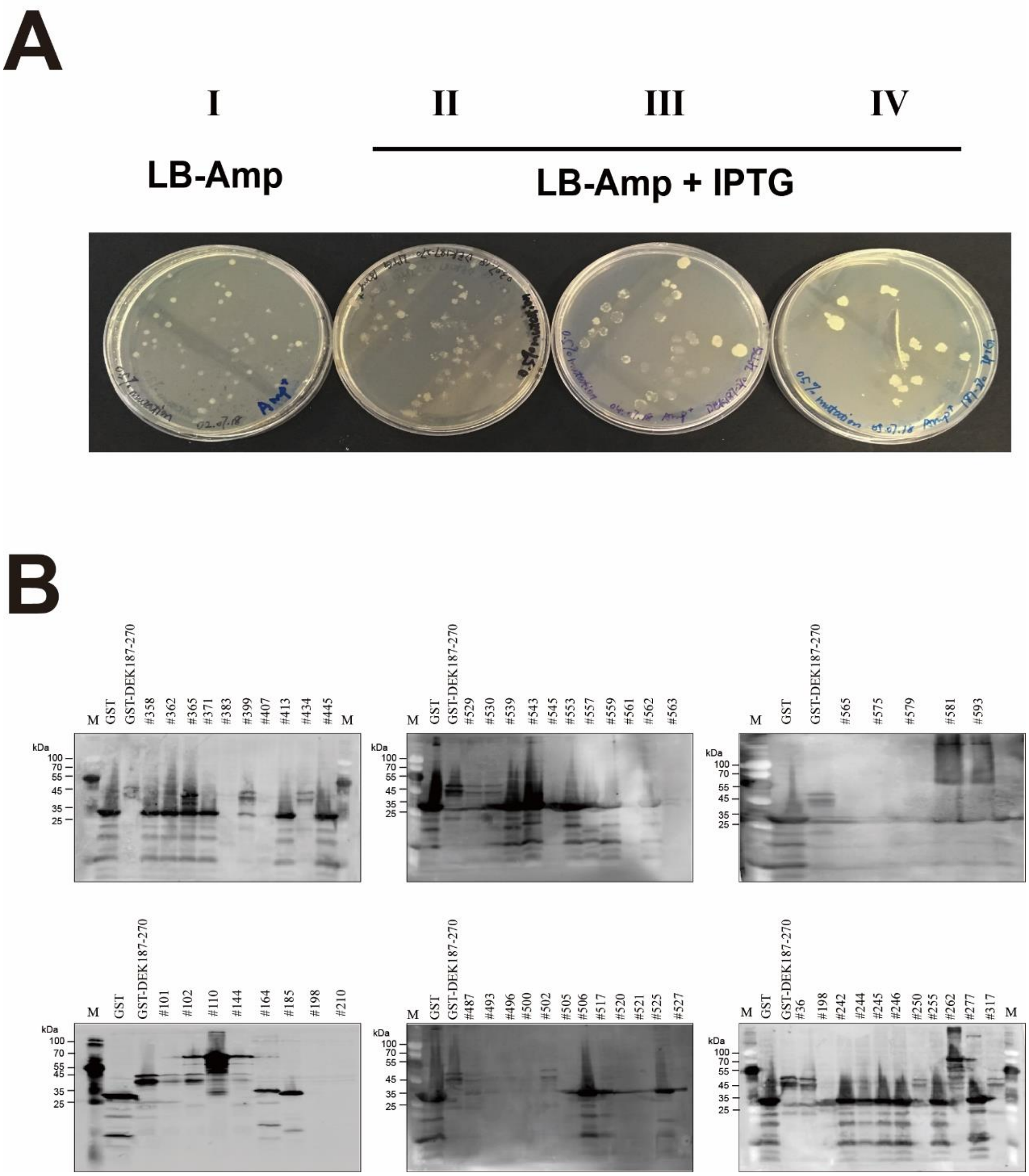


**Supplementary Figure 2. Bacterial Growth Inhibition Screen (BGIS): towards identification of a mutant in DEK 187-250 attenuated in the ability to bind to RNA.**

**A.** Mutants generated by error-prone PCR were ligated to pGEX-4T-1 plasmids, transformed, and initially plated on LB-Amp plates without IPTG (I). Subsequent replica stampings of the colonies to LB-Amp plates with IPTG were conducted to screen for mutants that lack bacterial growth inhibition (II-IV). Representative results from six screening rounds are shown.

**B.** Surviving colonies from (**A**) were subjected to small-scale expression, cell lysis and immunoblotting with antibodies specific to GST. GST only and GST-DEK187-270 served as negative and positive controls, respectively. Shown are blots from six rounds of mutational analysis.

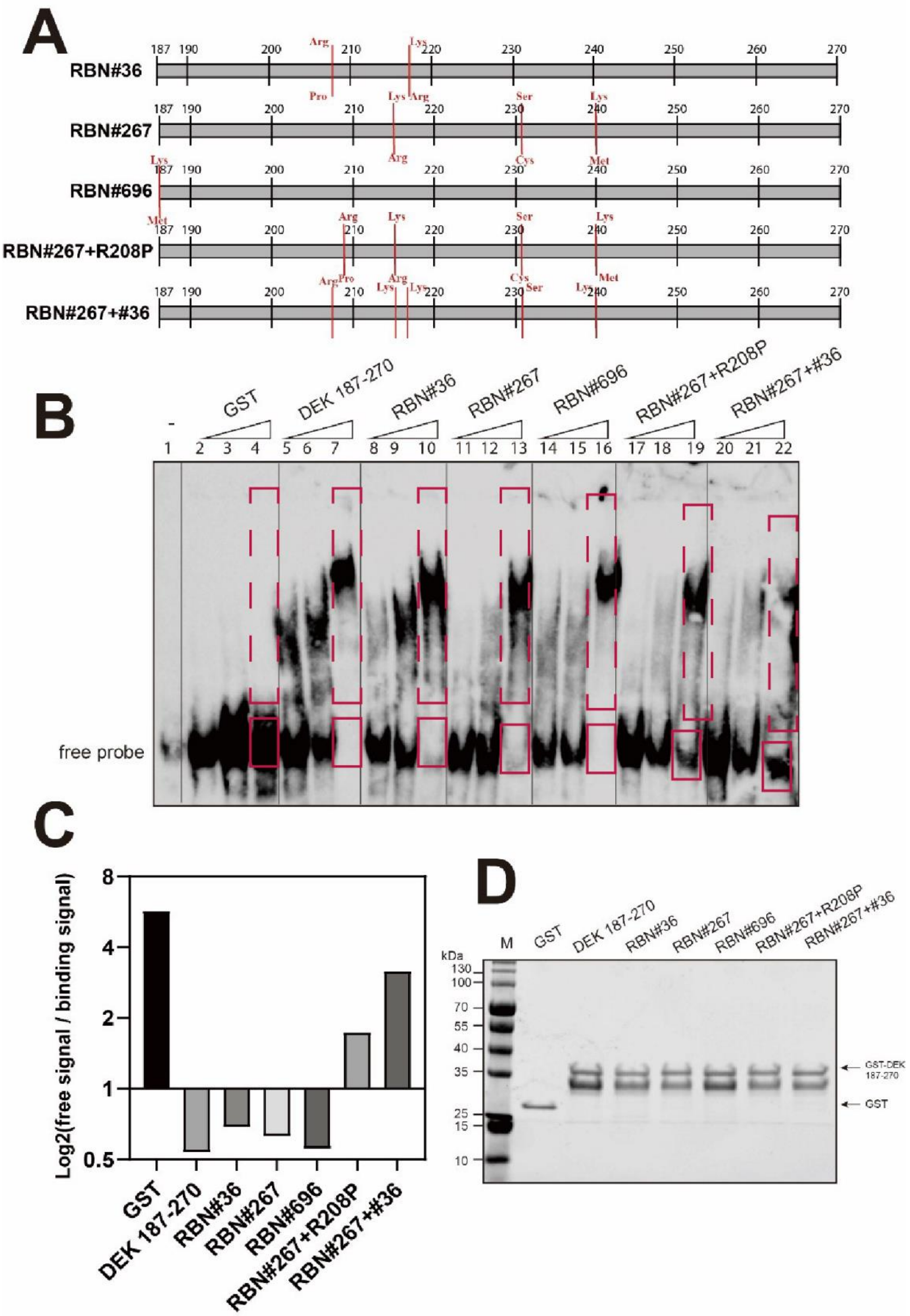

**Supplementary Figure 3. Evaluation of RNA binding affinities of BGIS-derived mutants or limited combinations of obtained mutations by RNA-EMSA**

**A**. Schematic depiction of mutation sites in each mutant with original amino acids on the top and identified mutation at the bottom;

**B**. RNA-EMSA was carried out as described in **Figure 1C** with GST-DEK fusions as indicated.

**C**. The grey value of the signals from free probe (solid red boxes) and the corresponding shifted bands (dashed red boxes) of the reactions with the highest protein amount was measured by ImageJ. The Log2 value of each free signal-to-binding signal ratio was calculated and Log2 of each value is shown in the bar chart, with the larger Log2 value indicating the weaker RNA binding affinity shown in (**B**);

**D**. Purified proteins used in each reaction in (B) were analyzed by SDS-PAGE and stained by Coomassie Blue to ensure equimolar input of the different protein preparations.

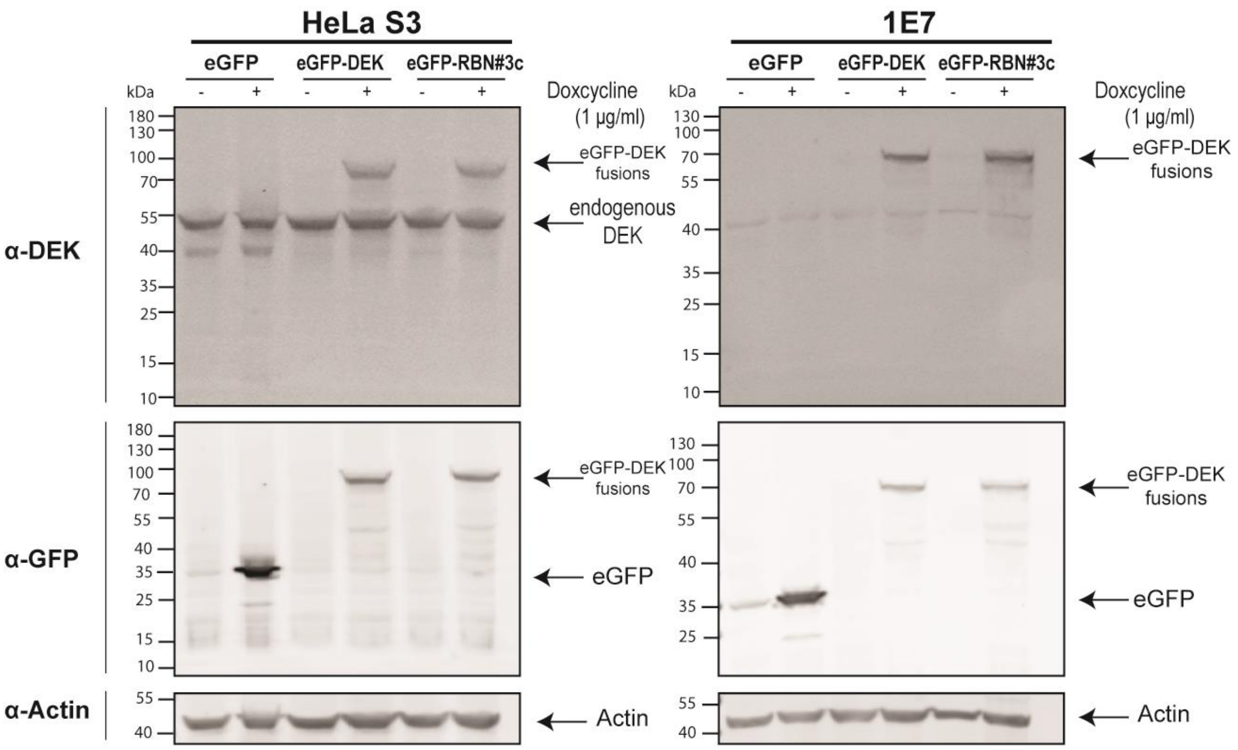


**Supplementary Figure 4: Analysis of stable, selected HeLa S3 and DEK KO (1E7) capable of inducible expression of eGFP-DEK fusions.**

Immunoblot analysis: HeLa or DEK KO (1E7) cell lines cultured on 10 cm cell culture dishes were treated with 1 µg/mL Doxycycline for 24 hours (+) or left untreated (-). Total cell lysates from $2x10^6$ cells were resolved by SDS-PAGE and subjected to immunoblot analysis with the indicated antibodies. Actin served as a loading control. Positions of molecular weight markers are indicated.

**A**

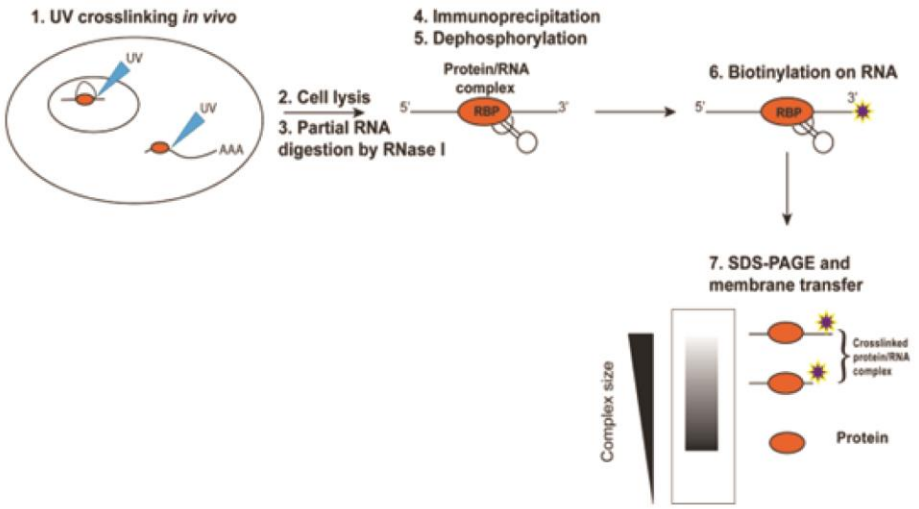


**B**

| Buffer Name | Modified CLIP buffer (Ule, Jensen, Mele, & Darnell, 2005)<br>I. | Modified iCLIP buffer (Huppertz et al., 2014)<br>II. | PAR-CLIP buffer (Danan, Manickavel, & Hafner, 2016)<br>III. | eCLIP buffer (E. Van Nostrand et al., 2016)<br>IV. | Modified PAR-CLIP (Bao et al., 2018)<br>V. |
|---|---|---|---|---|---|
| **Lysis buffer** | 50 mM Tris-HCl, pH 7.4<br>450 mM NaCl<br>0.5% NP-40<br>RNase inhibitor<br>Protease inhibitor cocktail | 50 mM Tris–HCl, pH 7.4<br>100 mM NaCl<br>1% Igepal CA-630<br>0.5% SDS<br>0.5% sodium deoxycholate<br>RNase inhibitor<br>Protease inhibitor cocktail | 50 mM HEPES, pH 7.5<br>150 mM KCl<br>2 mM EDTA<br>1 mM NaF<br>2% (v/v) NP40<br>0.5 mM DTT<br>Protease Inhibitor Cocktail<br>Phosphatase Inhibitor | 50 mM Tris-HCl pH 7.4<br>100 mM NaCl<br>1% NP-40<br>0.1% SDS<br>0.5% sodium deoxycholate<br>Protease Inhibitor Cocktail | 50 mM HEPES pH 7.5<br>150 mM KCl<br>2 mM EDTA<br>1 mM NaF<br>0.5% NP40<br>0.5 mM DTT<br>RNase inhibitor<br>Protease inhibitor cocktail |
| **High salt wash buffer** | 50 mM Tris-HCl, pH 7.4<br>750 mM NaCl<br>0.5% NP-40<br>0.1% SDS | 50 mM Tris-HCl, pH 7.4<br>1 M NaCl<br>1 mM EDTA<br>1% Igepal CA-630<br>0.1% SDS<br>0.5% sodium deoxycholate | 50 mM HEPES-KOH, pH 7.5<br>500 mM KCl<br>0.05% NP40 | 50 mM Tris-HCl pH 7.4<br>1 M NaCl<br>1 mM EDTA<br>1% NP-40<br>0.1% SDS<br>0.5% sodium deoxycholate | 50 mM HEPES-KOH pH 7.5<br>500 mM KCl<br>0.05% NP40<br>0.5 mM DTT<br>protease inhibitor cocktail |

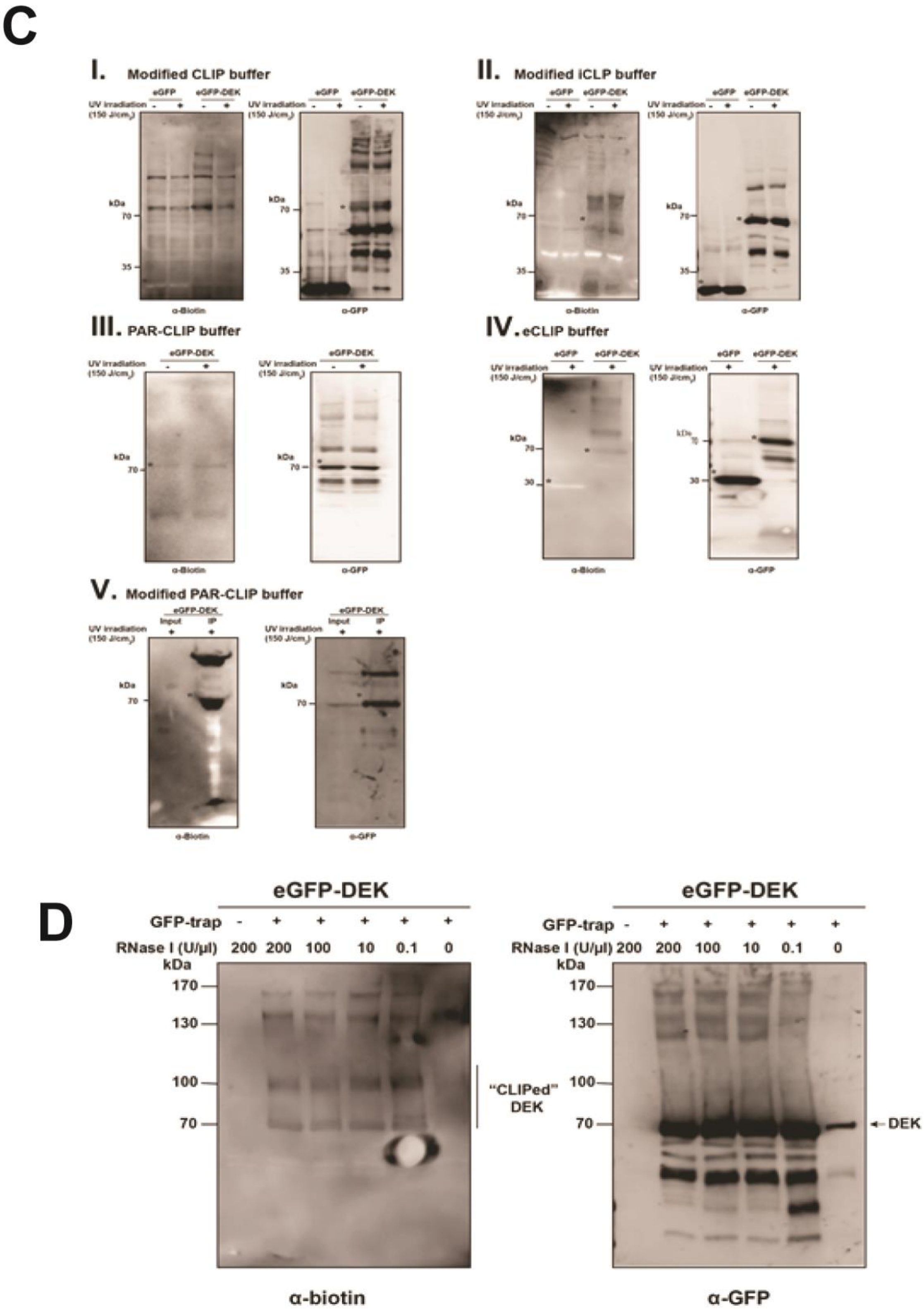


**Supplementary Figure 5: Optimization of CLIP for high affinity precipitations of DEK-RNA interactions.**

**A**. Schematic diagram of CLIP-seq applied in cells with doxycycline-inducible expression of eGPF tagged wt or RBN#3c DEK. Cells, after induction of protein expression, are crosslinked by UV irradiation, followed by RNase I fragmentation (1-3). GFP-trap was added to immunoprecipitate GFP tagged DEK and crosslinked RNAs (4). After stringent wash steps, RNA fragments are dephosphorylated to enable a biotinylated cytidine (bis)phosphate to ligate to 3' end of each strand (5 and 6). The mixture is then resolved by SDS-PAGE and transferred to a nitrocellulose membrane for detection (7). Relevant complexes, as indicated by brackets, on the membrane are extracted, subjected to proteinase K treatment and purification for subsequent high throughput RNA sequencing (8 and 9).

**B**. CLIP buffer compositions derived from indicated protocols [4-8] and used for CLIP optimization in this study.

**C**. Stringencies of buffers are crucial and need to be optimized according to every protein of interest. Five CLIP buffer systems, as listed in the table above, were tested for their suitability in our chosen cellular system overexpressing eGFP-DEK. After CLIP immunoprecipitation, half of the IP samples along with corresponding inputs were loaded on a HEPES gel and processed for detection of biotin (left panels of each buffer system) and the remaining samples were subjected to regular immunoblotting with GFP-specific antibodies (right panels). The asterisks indicate the positions of eGFP or the eGFP-DEK fusion corresponding to their regular (non-crosslinked) expected migration position.

**D.** Optimization of ribonuclease treatment in CLIP. Left panel: Detection of biotin labelled RNA-DEK complexes after immunoprecipitation from lysates treated with different concentrations of RNase I, ranging from 0 to 200 units per microliter. The region indicated as “CLIPed” DEK was determined as RNA fragments of desired length. Right panel: Immunoblot analysis of samples as on the left using GFP-specific antibodies. The expected molecular weight of GPF-DEK (approximately 70 kDa) and the immunoprecipitation efficiency of GFP-trap among the lysates with various concentrations of RNase I is shown. Molecular weight markers are indicated.

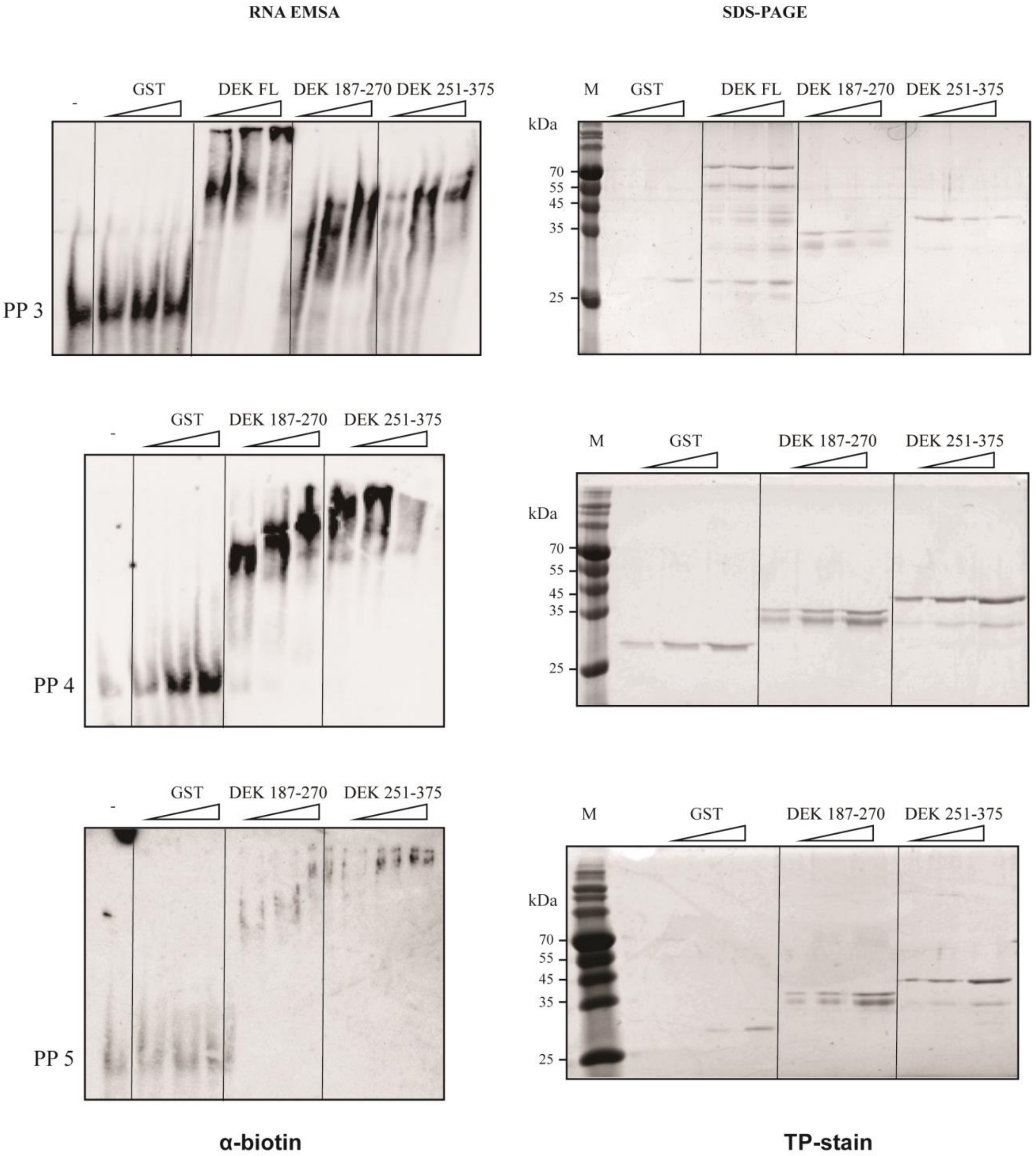


**Supplementary Figure 6: RNA-EMSA indicates two RNA interaction domains in DEK.**

Left panels: single stranded RNA pentaprobes (PP3-PP5) were incubated with increasing amounts of GST tagged DEK fragments (DEK 187-270 and DEK 251-375), GST only or left untreated (-). RNA-EMSA was carried out as described in Figure 1C.

Right panels: purified proteins used in each EMSA reaction were analysed by SDS-PAGE and stained by Coomassie Blue to ensure equimolar input of the different protein preparations.

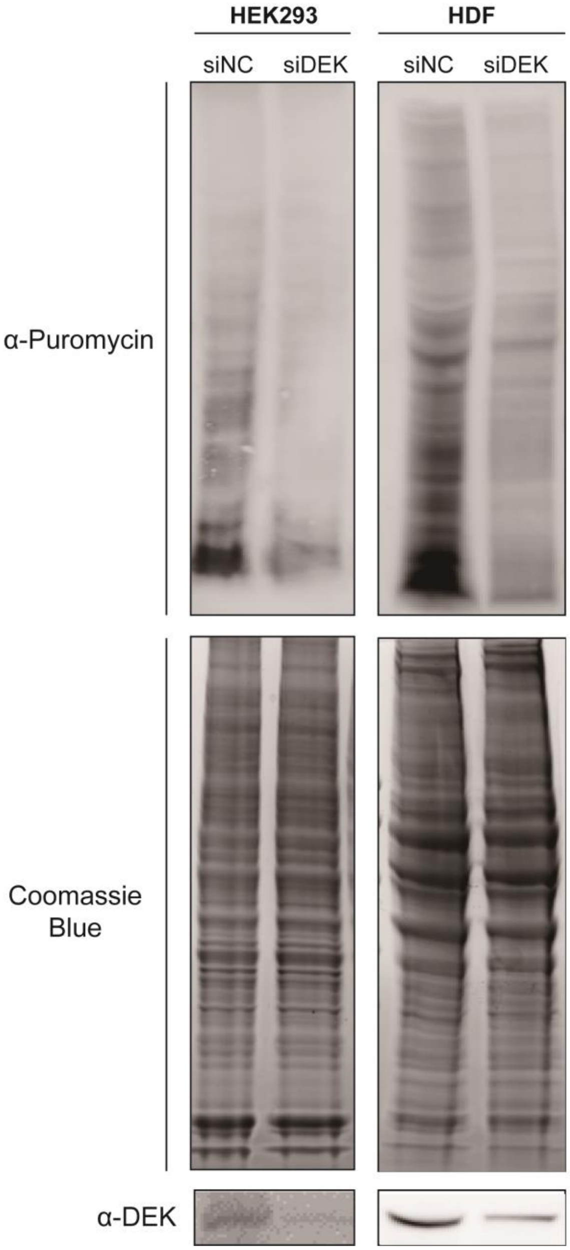


**Supplementary Figure 7. SUnSET assay monitoring nascent protein synthesis rate upon DEK depletion shows reduced new protein synthesis in HEK293 cells and in primary human dermal fibroblasts.**

Protein translation was monitored after treatment of cells with puromycin at a final concentration of 10 μg/mL for 10 minutes as described in Figure 3. HEK293 or HDF cells transfected with siDEK or siNC were labelled with puromycin followed by total protein extraction. Half of the extract was separated by SDS-PAGE and analysed by immunoblotting with antibodies specific to puromycin (12D10) (blot on top) or DEK (blot at bottom), and another half was analysed by SDS-PAGE and stained by Coomassie Blue to monitor total protein loading.

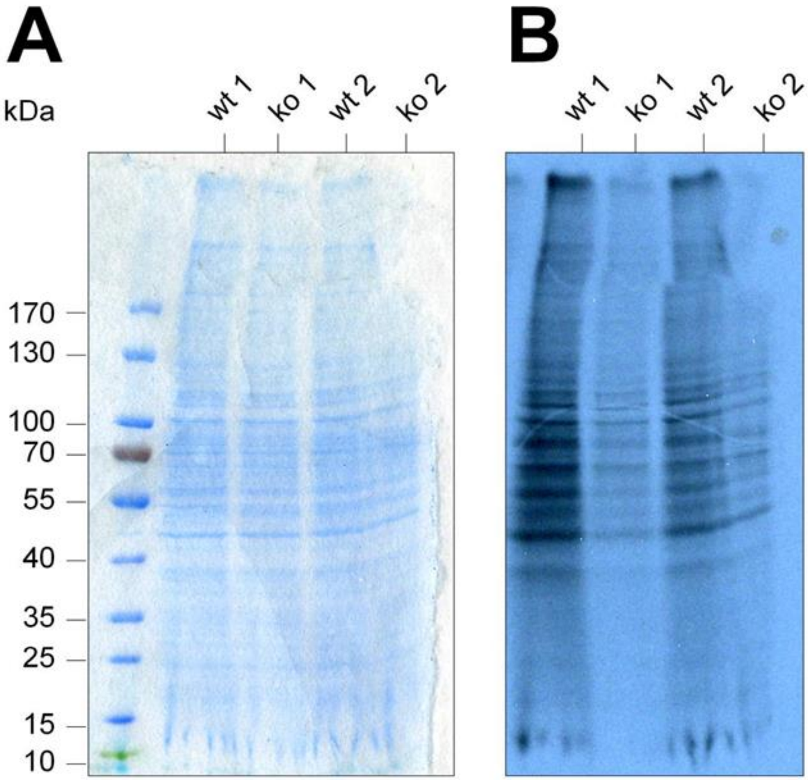


**Supplementary Figure 8: DEK knockout cells show reduced $^{35}$S methionine incorporation in HeLa S3 cells.**
HeLa S3 control (wt) and DEK knockout (KO) cells were seeded in six-well plates one day prior labelling. Cells were labelled with 50 µCi/well $^{35}$S-methionine in methionine free medium for 2 h. The cells were washed, lysed in lysis buffer and separated via SDS-PAGE and the gel was dried.

**A**. Total protein stain using Coomassie. Size markers are indicated on the left (kDa).

**B**. The dried gel from (**A**) was exposed to an X-ray film.

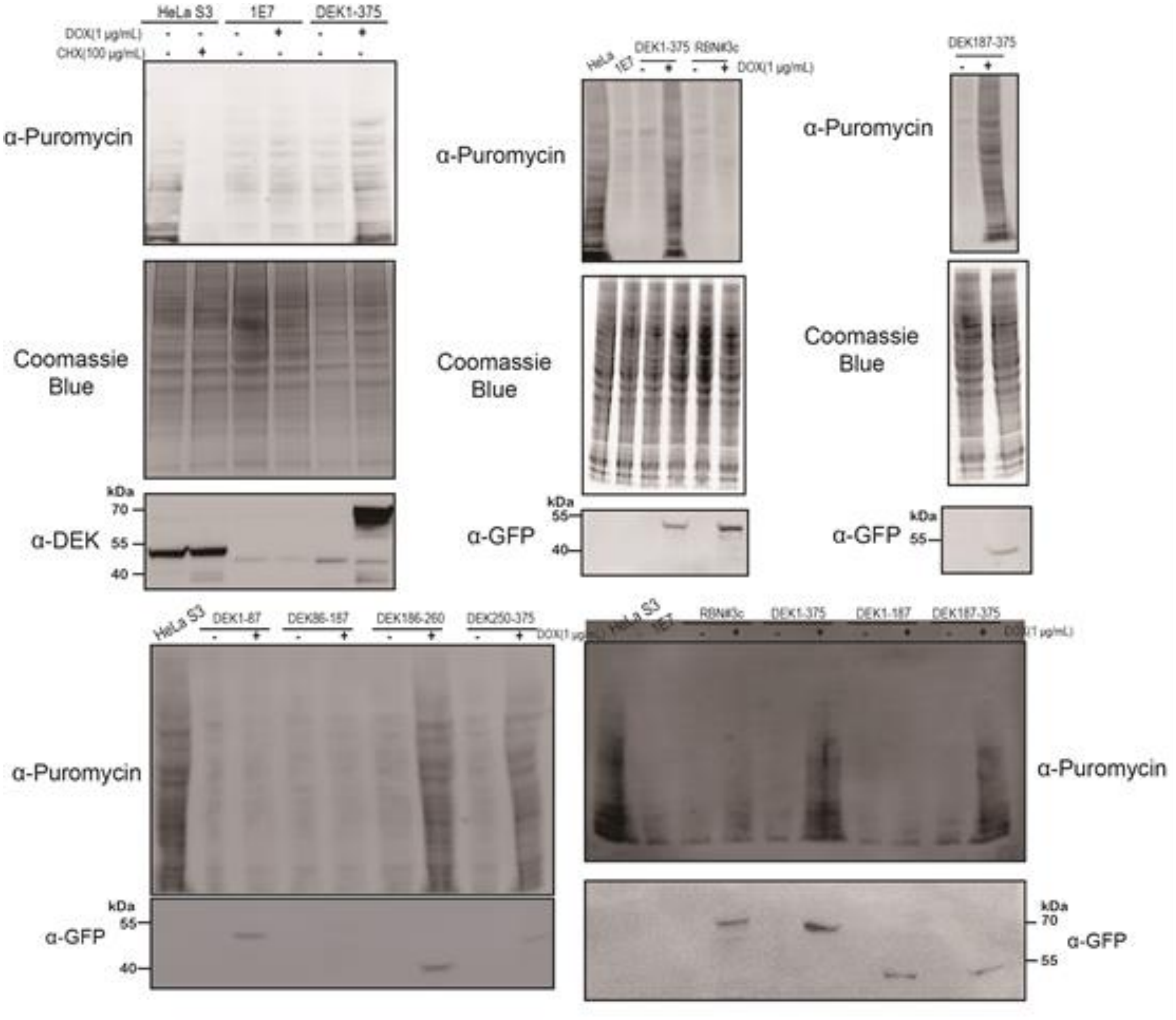


**Supplementary Figure 9: SUnSET assay, monitoring nascent protein synthesis rate, reveals functional implications of the DEK-C terminal domain in the regulation of ribosome function (related to Figure 3 A and B).**

Shown are repetitions of experiments as outlined in Figure 3.A and B. Cells either treated with CHX with indicated concentration for 10 minutes at RT or treated with DOX with indicated concentration for 24 hours were followed with treatment with puromycin solution to a final concentration of 10 µg/mL for 10 minutes for the purpose of nascent proteins labelling. After cell collection and concentration determination, the extracts were separated by SDS-PAGE followed by immunoblotting with antibodies specific to puromycin (12D10), GFP or DEK. A fraction of the extracts was stained by Coomassie Blue to ensure equal amounts of proteins loaded in each lane.

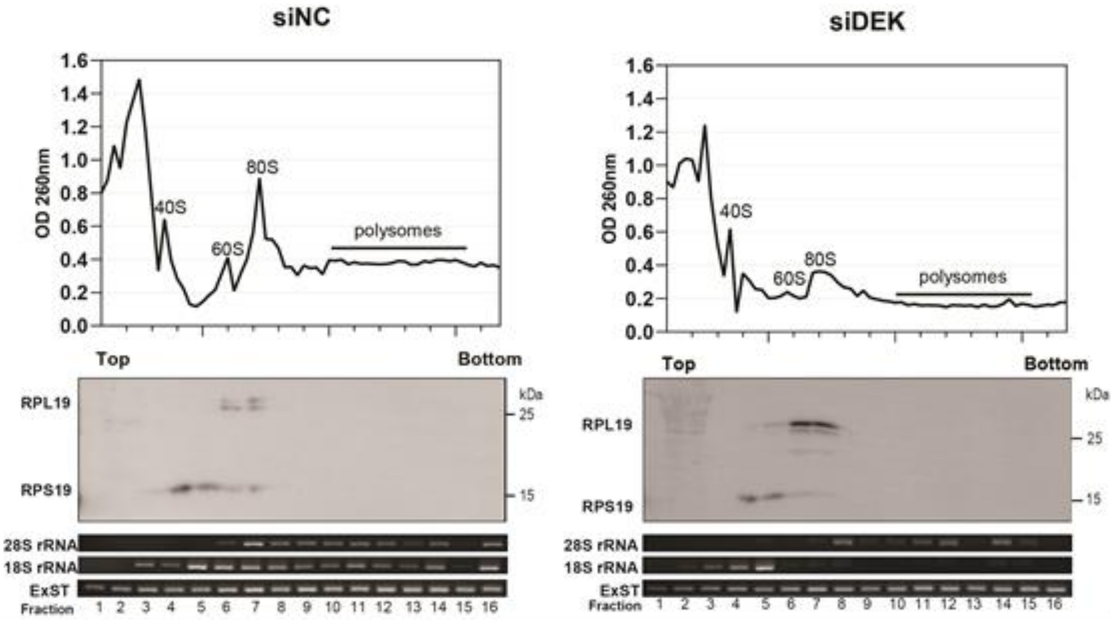


**Supplementary Figure 10: Knockdown of DEK expression induces alterations to quantity and quality of ribosome in HeLa S3 cells (related to Figure 3C).**
Shown is an additional biological replicate of cytoplasmic ribosome profiling, which was carried out as described in Figure 3C. (**Upper panels**). An aliquot of each fraction was purified and analysed by SDS-PAGE and immunoblotting with antibodies specific to RPS19 and RPL19 to identify the positions of 40S, 60S and 80S ribosomes; Similarly, positions of 18S and 28S rRNA were also analysed by PCR after RNA extraction and reverse transcription; ExST: External Standard (**Lower panels**).

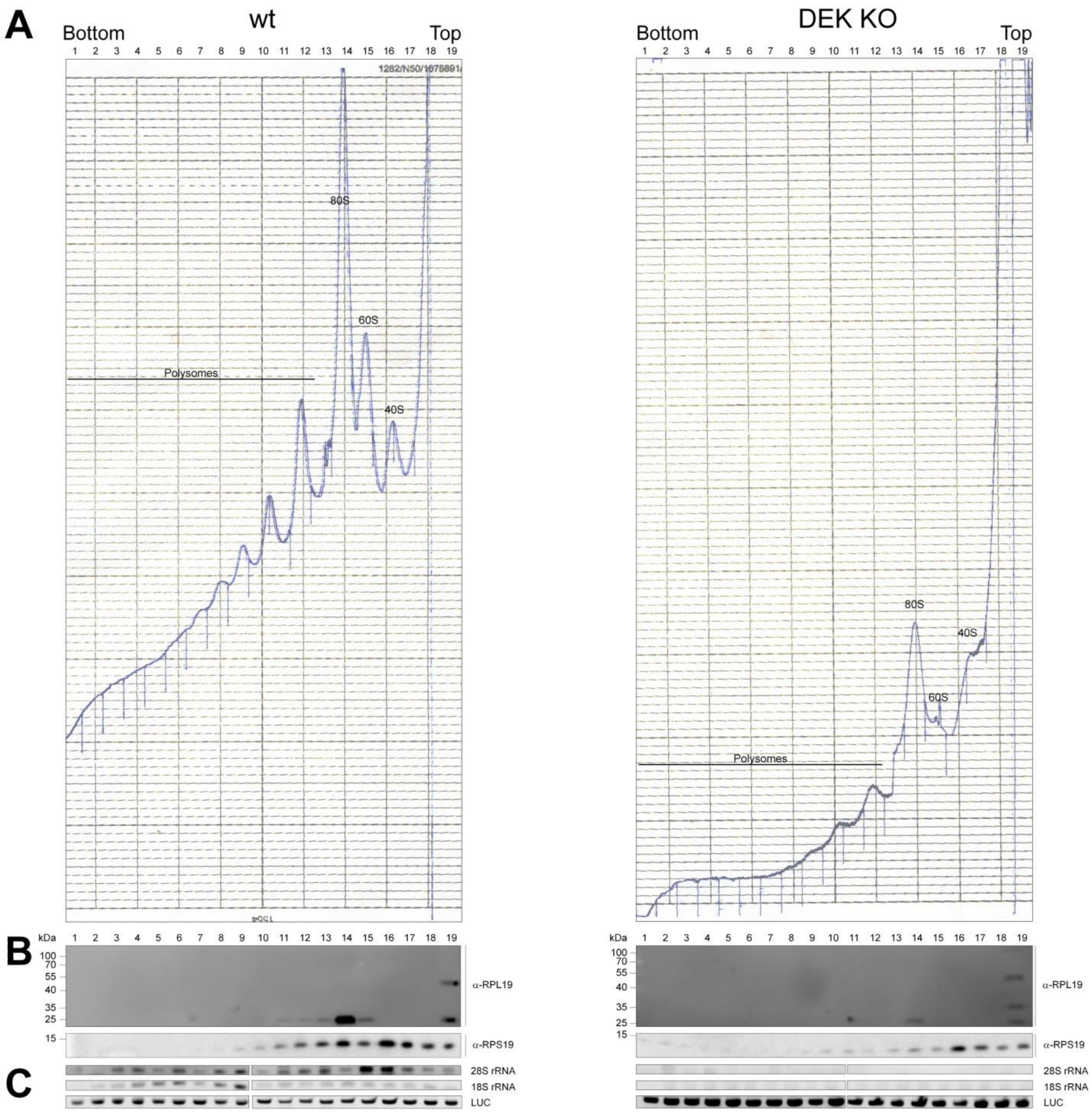


**Supplementary Figure 11: Depletion of full length DEK induces altered quantity and quality of ribosomes in cytoplasmic extracts (additional biological replicate performed under different experimental settings).**

4 x $10^6$ HeLa S3 and DEK knockout cells were seeded two days prior treatment. After two days the cells were incubated with 100 µg/mL cycloheximide for 10 min at 37°C and cells were washed three times with ice-cold PBS containing cycloheximide. Cells were harvested by scraping, dounced, and cell debris and nuclei were removed by centrifugation at 13,000 g at 4°C for 10 min.

**A**. 800 µg of cytoplasmic extract from wt HeLa S3 wt and DEK knockout cells were layered on a 15 – 45 % sucrose density gradient in the presence of 100 µg / mL cycloheximide. Density separation was carried out via ultracentrifugation (2 h, 45,000 rpm, SW41Ti Rotor) and 20 fractions were collected using a Frac-920 from GE Healthcare. During fractionation presence of nucleic acids was monitored at 260 nm with an UVis-920 from GE Healthcare device and recorded on a Rec111 (GE Healthcare) writer. Based on the OD260 nm profile the positions of polysomes, the complete ribosome 80S, and the large 60S or small 40S subunits are indicated within the graph.

**B**. Analysis of selected ribosomal proteins: Proteins from each fraction were precipitated and analyzed via SDS-PAGE with subsequent immunoblotting using RPL19- and RPS19-specific antibodies. Size markers are indicated on the left (kDa).

***C***. Analysis of rRNA: First, identical amounts of LUC RNA were spiked into samples as in B, followed by RNA extraction, reverse transcription and analysis via PCR using specific primer pairs for the 28S, 18S rRNA forms and for Luc, which served as an internal control for RNA purification efficiency.

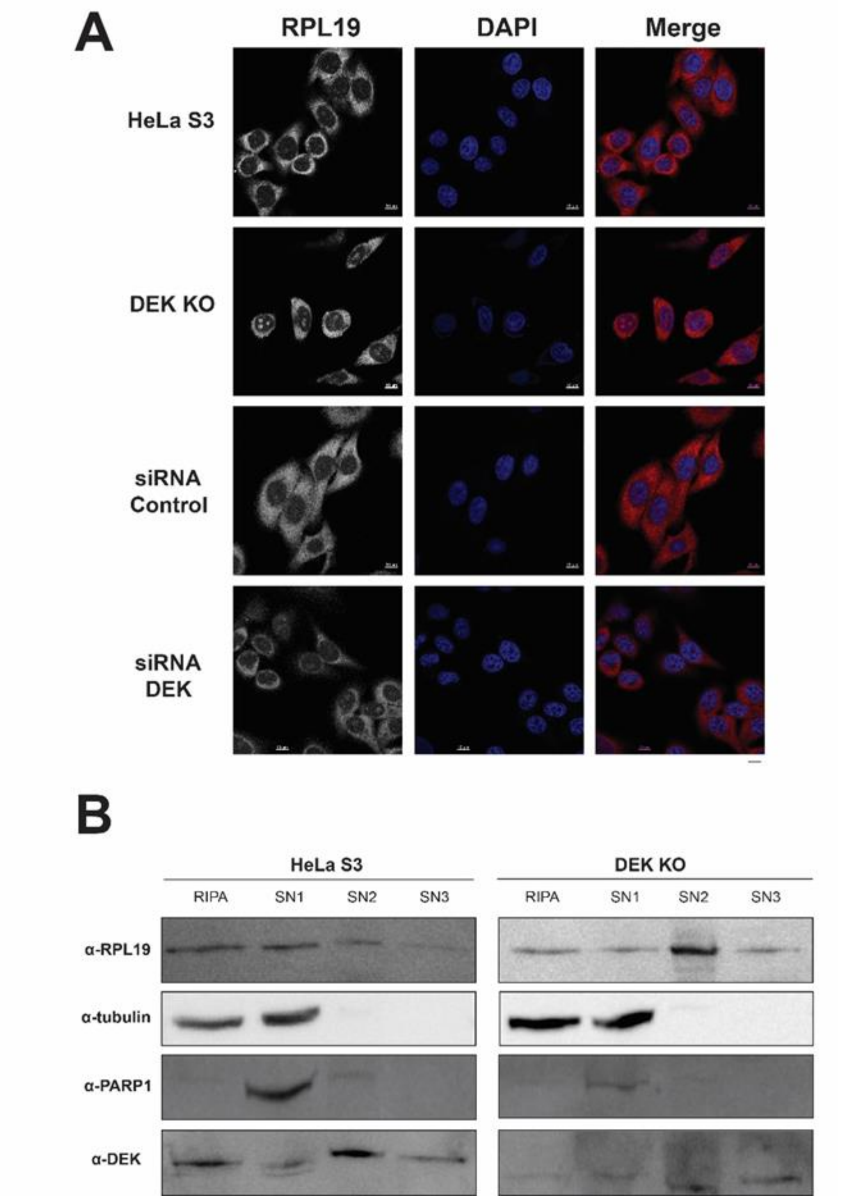


**Supplementary Figure 12*:* Depletion of DEK induces alterations to the subcellular localization of RPL19.**

**A.** HeLa S3 and DEK KO cells or HeLa S3 cells transfected with siRNA targeting *DEK* or nonsense sequence were seeded on pre-coated glass slides one day before treatment, followed by fixation, permeabilization and hybridization with antibodies specific to RPL19. DNA was stained with DAPI. Confocal images were captured with a Zeiss LSM880. Scale bar: 10 μm. Shown are representative micrographs from three independent experiments.

**B.** Pre-ribosomal complexes were isolated by sequential extraction method according to their experimentally defined accessibility. The extracts from resulting fractions, namely SN1, SN2 and SN3, along with one-step extraction using RIPA buffer were loaded on SDS-PAGE gels followed by immunoblotting with indicated antibodies specific to DEK and RPL19, anti-tubulin and PARP1 were used as internal controls. Shown are the representative immunoblotting results from two independent experiments.

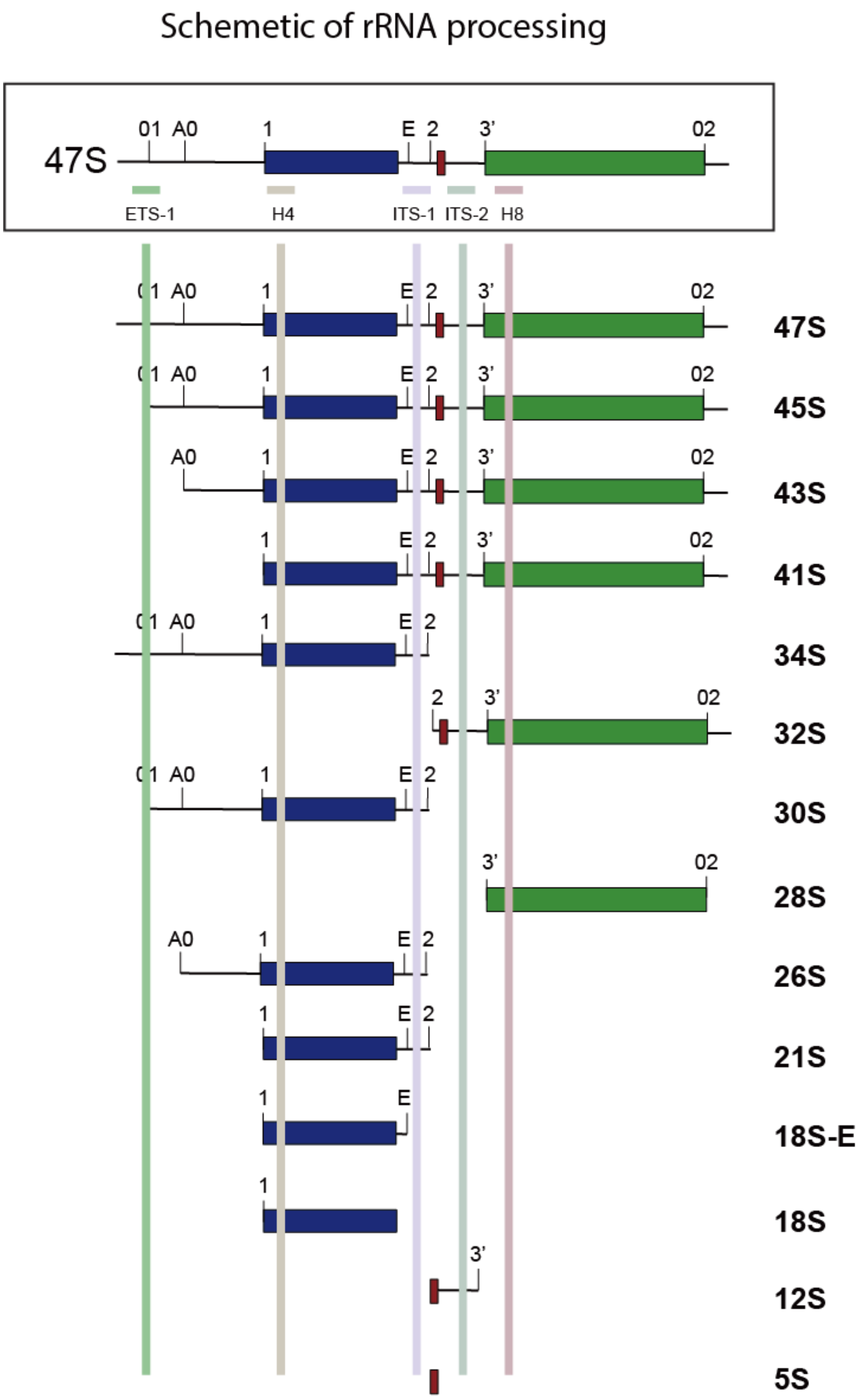


**Supplementary Figure 13: Schematic overview of the major 47S pre-rRNA processing pathway in HeLa cells with the location of probes designed for qPCR indicated**.

In HeLa cells, the maturation process starts from the generation of 18S rRNA by exonucleolytic and endonucleolytic cleavages at 5’ETS and ITS1 followed by trimming of flanking sequences of 5.8S, and then the removal of ITS2 spacer sequence is initiated to allow the maturation of 28S and 5.8S rRNA.

See **Supplementary Tables 5-7** for probe sequences.

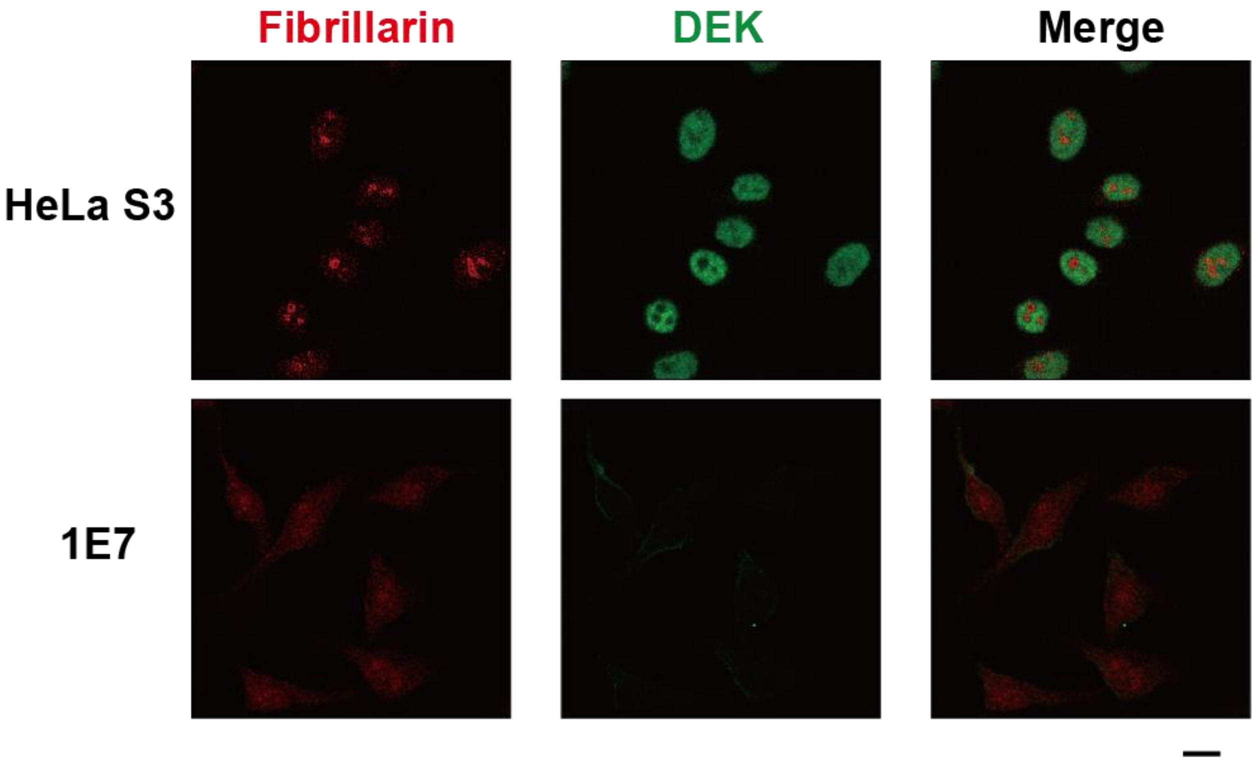


**Supplementary Figure 14: Alterations to the morphology of nucleoli upon depletion of DEK.**

HeLa S3 and DEK KO cells were seeded on pre-coated glass slides one day before treatment, followed by fixation, permeabilization and hybridization with antibodies specific to fibrillarin, a nucleolar marker, and monoclonal antibodies specific to DEK. DNA was stained with DAPI. Images were captured with a Zeiss LSM880 confocal microscope. Scale bar: 10 μm.

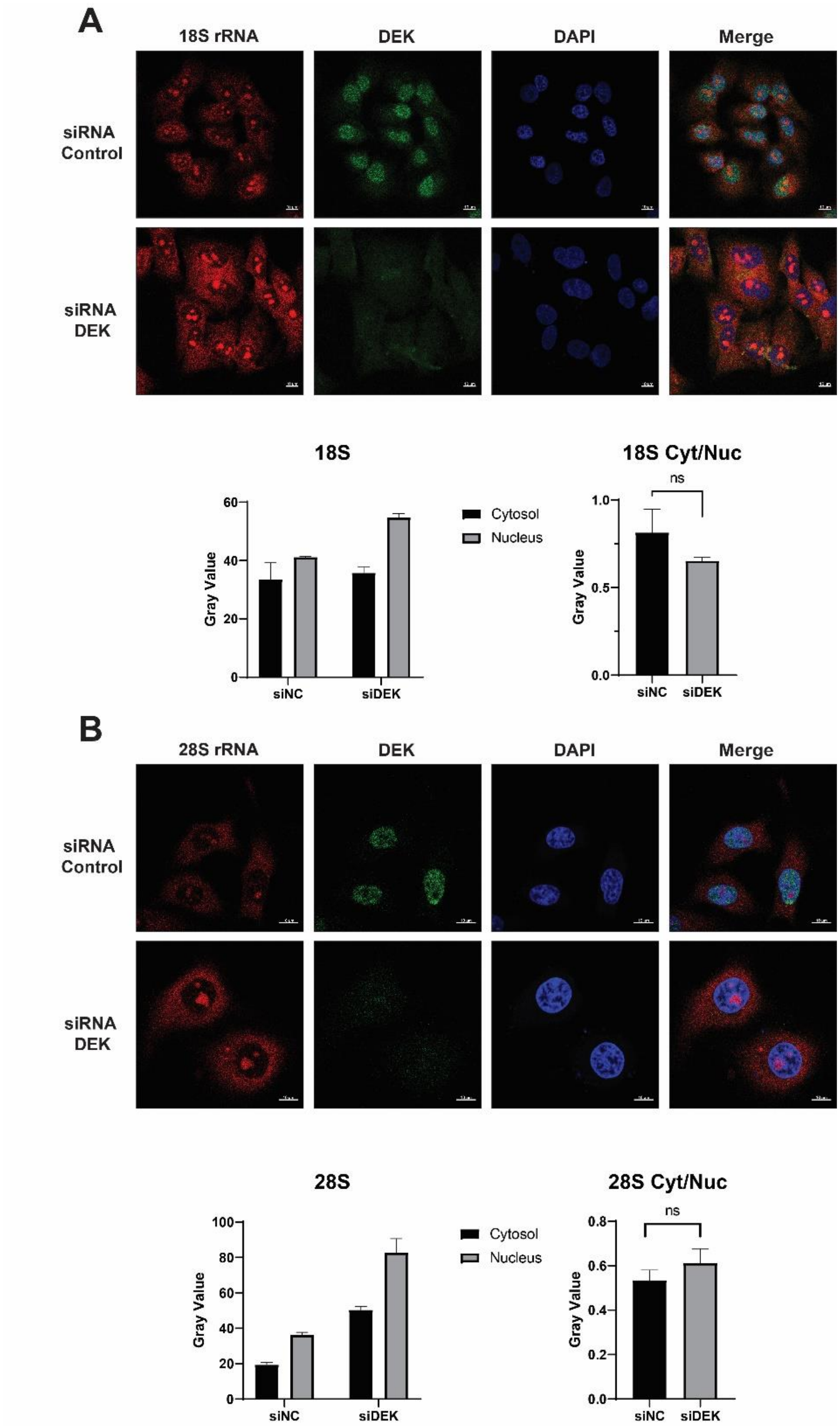
A
18S rRNA
DEK
DAPI
Merge
siRNA Control
siRNA DEK
18S
Gray Value
60
40
20
0
siNC
siDEK
Cytosol
Nucleus
18S Cyt/Nuc
ns
1.0
0.5
0.0
B
28S rRNA
DEK
DAPI
Merge
siRNA Control
siRNA DEK
28S
100
80
60
40
20
0
Cytosol
Nucleus
28S Cyt/Nuc
ns
0.8
0.6
0.4
0.2
0.0

**Supplementary Figure 15: Subcellular distribution of 18 S and 28S rRNA upon depletion of DEK.**

HeLa S3 cells transfected with siRNAs targeting DEK or a nonsense sequence were seeded on coated glass slides one day before treatment. They were fixed, permeabilized and hybridized with Cy5 labelled probes annealing to 18S (**A**) or 28S (**B**) rRNA, followed by immunostaining with monoclonal antibodies specific to DEK and DNA was stained with DAPI (blue). Confocal images were taken by a Zeiss LSM880 microscope**.** Scale bar: 10 µm; For quantitative analysis, the DAPI channel was recognized by the macro of ImageJ as the nuclei. The threshold-mask was then adjusted to 40 to separate signals of the cytoplasmic 18S/28S rRNA from background. Average pixel intensity and pixel coverage were measured giving rise to relative intensities from cytoplasm and nucleus. The ratios of cytoplasmic to nucleic intensities were calculated with approx. Forty cells (n=40) were taken from four images for each measurement. Student's t-test (Two-tailed) was applied to the results.

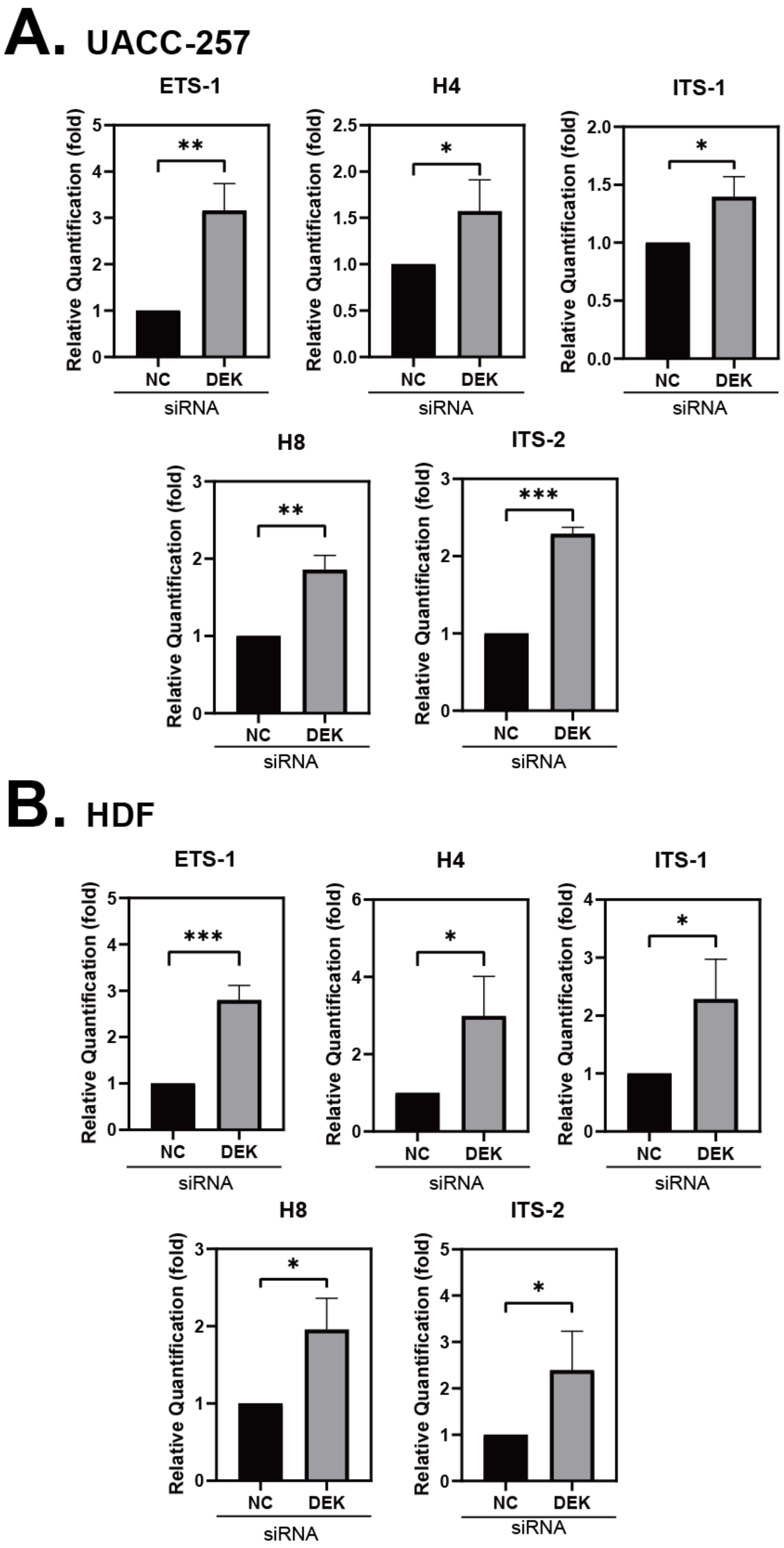
A. UACC-257
ETS-1
H4
ITS-1
H8
ITS-2
Relative Quantification (fold)
NC
DEK
siRNA
B. HDF
ETS-1
H4
ITS-1
H8
ITS-2
Relative Quantification (fold)
NC
DEK
siRNA

**Supplementary Figure 16: DEK depletion results in altered expression of various pre-rRNA species.** Quantitative analysis of indicated rRNA species by qPCR in melanoma cells (UACC-257) (**A**) and primary human dermal fibroblasts (HDF) cells (**B**) transfected with siRNAs targeting DEK or with nonsense siRNAs using the primers indicated in **Supplementary Figure 13**. Student's t-test (Two-tailed) was applied to three biological repliactes. Asterisks indicate the according p-value (*$p < 0.05$, **$p < 0.01$, ***$p < 0.005$).

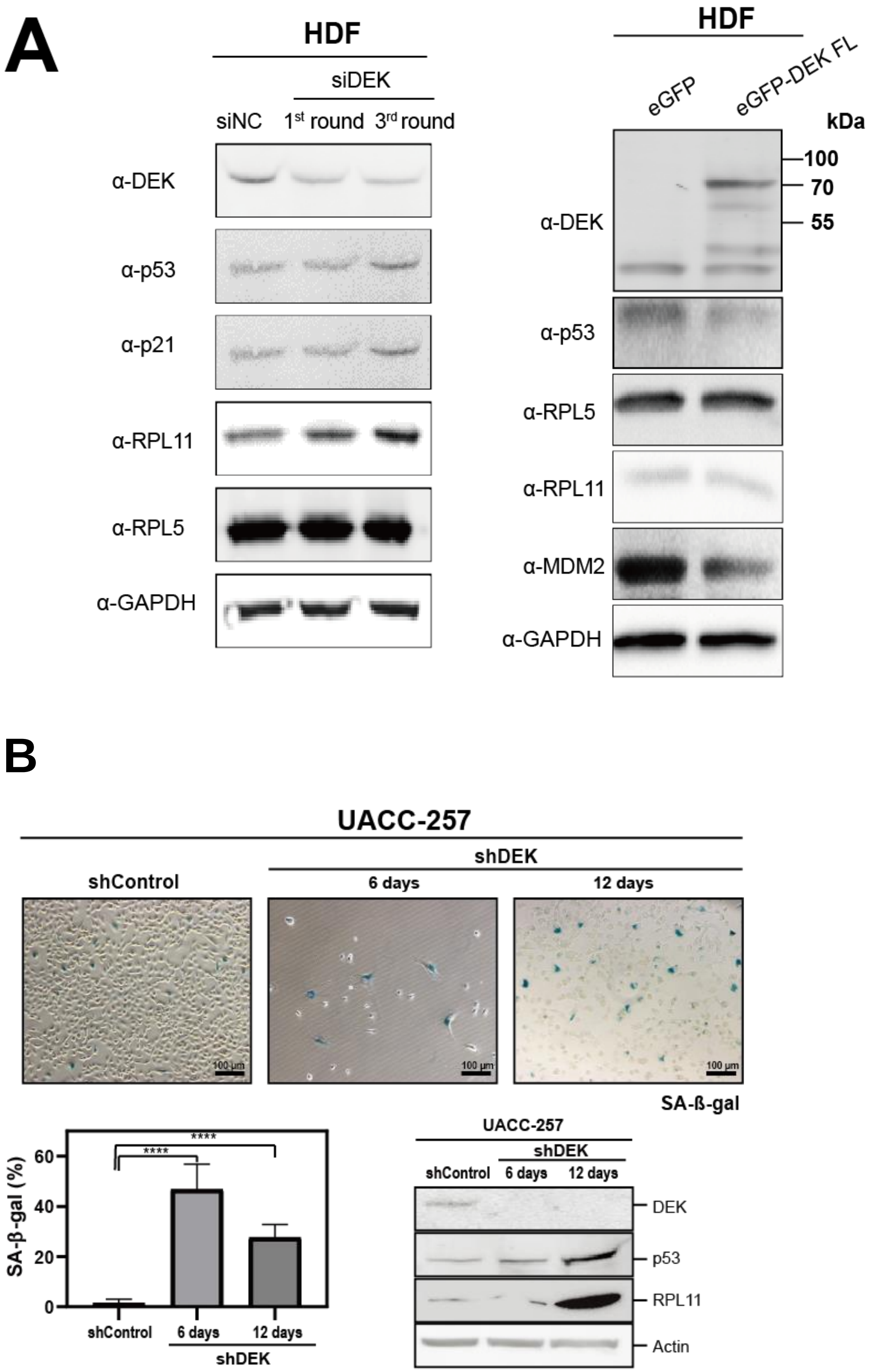
A
HDF
siDEK
siNC
1st round
3rd round
α-DEK
α-p53
α-p21
α-RPL11
α-RPL5
α-GAPDH
HDF
eGFP
eGFP-DEK FL
kDa
100
70
55
α-DEK
α-p53
α-RPL5
α-RPL11
α-MDM2
α-GAPDH
B
UACC-257
shControl
shDEK
6 days
12 days
100 μm
SA-ß-gal
SA-β-gal (%)
60
40
20
0
****
****
shControl
6 days
12 days
shDEK
UACC-257
shDEK
shControl
6 days
12 days
DEK
p53
RPL11
Actin

**Supplementary Figure 17*:* Evaluation of expression level of key proteins involved in IRBC and induction of cellular senescence induced by depletion of DEK.**

**A.** **Left panel**: Lysates of HDF cells post one round or three rounds of transfections with siRNAs targeting DEK along with the control were analysed by immunoblotting with antibodies specific to key proteins in the IRBC pathway, including RPL5, RPL11, p53 and p53 targeting protein p21. GAPDH served as an internal control; **Right panel**: Lysates of HDF cells with overexpression of eGFP-DEK FL and eGFP only as control were analysed by immunoblotting using the indicated antibodies.

**B.** Metastatic melanoma (UACC257) cells were transduced with lentiviral particles either delivering plasmids containing shRNA targeting DEK or a scrambled shRNA. shDEK cells were stained with SA-β-gal 6 days and 12 days post of transductions, as well as the control cells 12 days post transduction. Images were captured by Nikon Ti-S inverted fluorescence microscopy. Shown are the representative images of each cell type. Scale bar: 10 μm; Four images of each cell type were analysed and the ratio of senescent cells (in blue) to total cells were calculated. Student's t-test (Two-tailed) was applied to the result. Asterisks indicate the according p-value (****$p < 0.001$). Cell lysates on day 6 and day 12 after infection along with the control on day 12 were analysed by immunoblotting with the indicated antibodies.

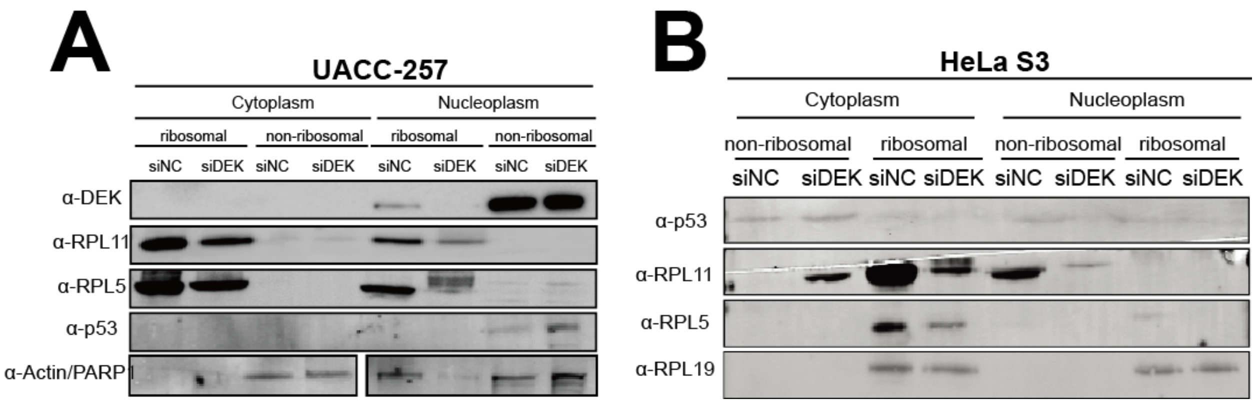


**Supplementary Figure 18: Redistribution of cellular RPL5 and RPL11 proteins upon DEK depletion.**

**A.** Melanoma cells (UACC-257) were transfected with siRNA targeting DEK or nonsense siRNA for 72 hours. Ribosomal and non-ribosomal fractions of cytoplasmic and nucleoplasmic extracts were isolated by ultracentrifugation. The same amount of total protein as determined by BCA for each fraction was analysed by immunoblotting with the indicated antibodies. Actin was used as a cytoplasmic control and PARP1 was served as a nucleoplasmic control.

**B**. HeLa S3 cells were transfected with siRNAs targeting DEK or a negative control siRNA for 72 hours. Ribosomal and non-ribosomal fractions of cytoplasmic and nucleoplasmic extracts were isolated by ultracentrifugation. The same amounts of proteins as determined by BCA for each fraction were analysed by immunoblotting with the indicated antibodies.

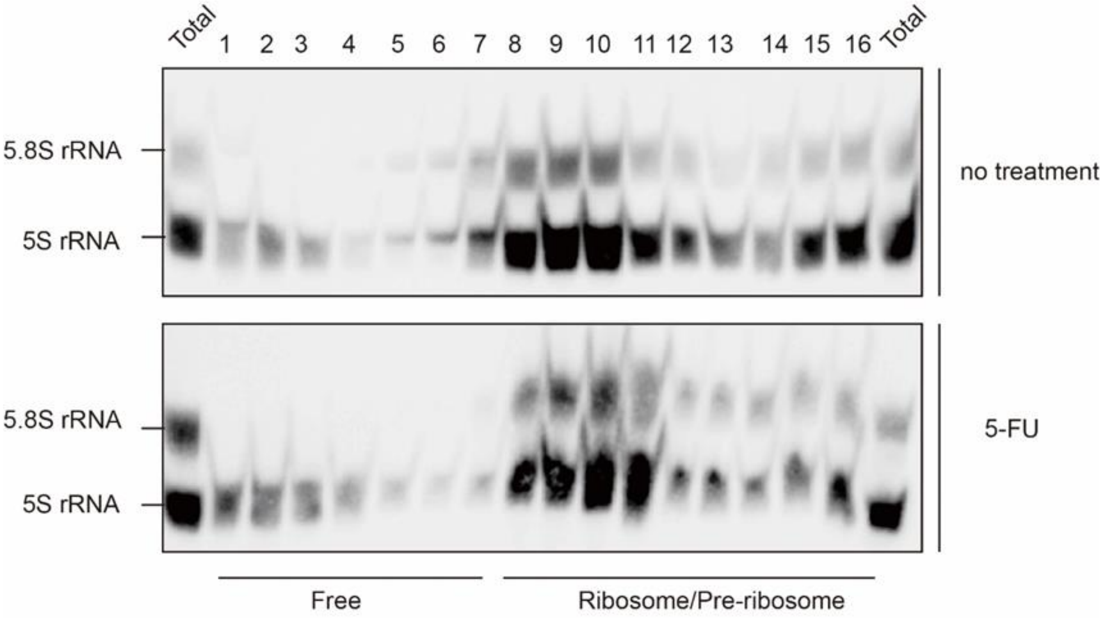


**Supplementary Figure 19: 5-FU treatment leads to redistribution of 5S rRNA.**

Total RNA of HeLa S3 cells treated with 5-FU at a final concentration of 100 μM or without treatment was separated by polyacrylamide gel electrophoresis and analysed by Northern Blot with probes annealing to 5S rRNA and 5.8S rRNA, which served as a control.

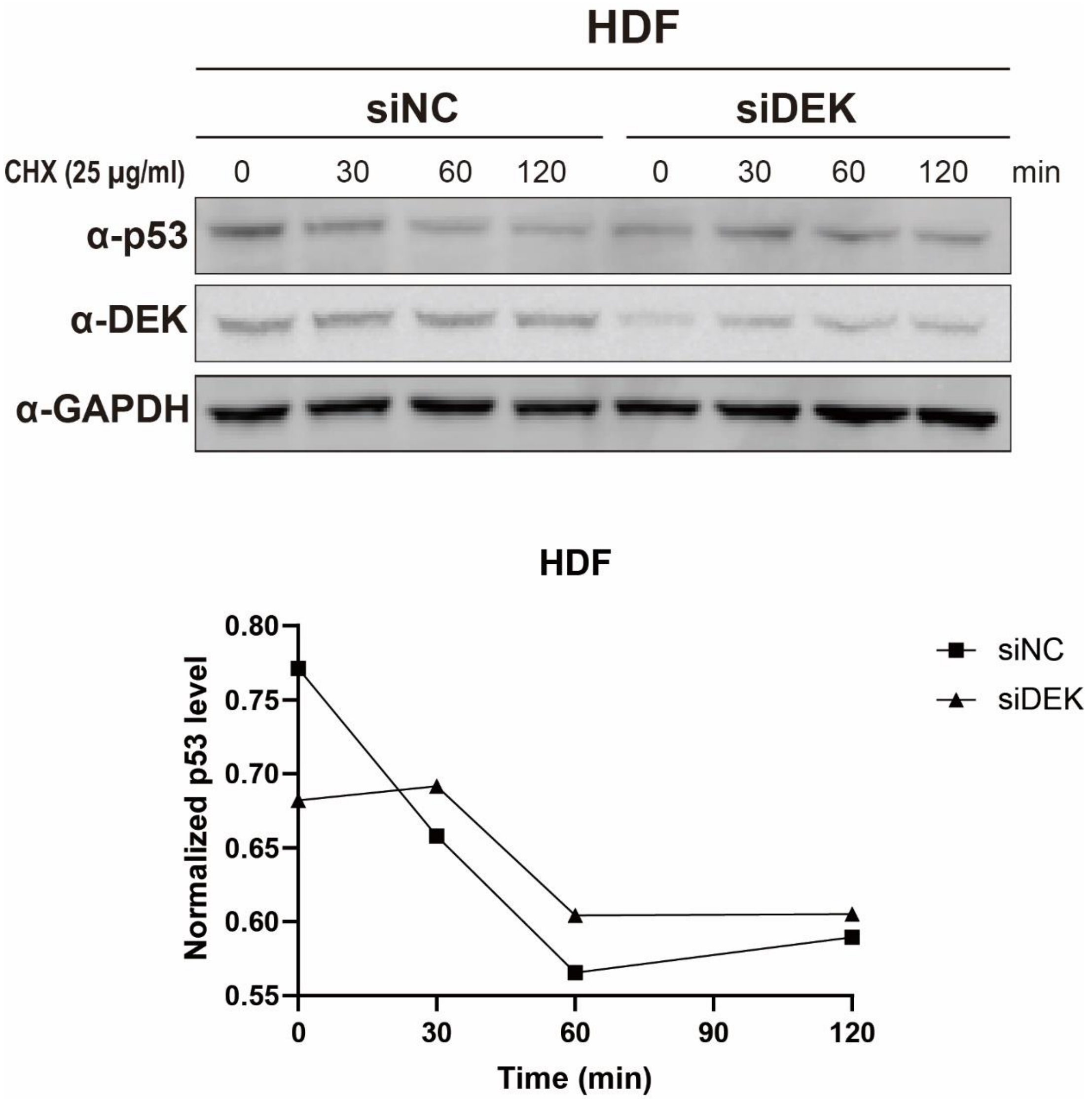


**Supplementary Figure 20: DEK regulates P53 stabilization (biological replicate of Figure 6E).**

Time-lapse examination of p53 degradation dependent on DEK. Human dermal fibroblast (HDF) primary cells transfected with siRNA targeting DEK or a negative control siRNA for 72 hours were seeded one day before treatment. Cells were treated with 25 µg/mL cycloheximide (CHX) and collected at the indicated time points and analysed by immunoblotting with the indicated antibodies. The signal intensities were measured by ImageJ and plotted as the intensities of p53 normalized to GAPDH. Shown is a repetition of the experiment as outlined in Figure 6E.

### Supplementary Tables 1-8

##### Supplementary Table 1: Top ten GO terms enriched in DEK wt CLIP peaks generated by DAVID analysis

| Category | log2FoldChange | -log10FDR |
|---|---|---|
| GO:0006413~translational initiation | 1.060792 | 24.75336 |
| GO:0022625~cytosolic large ribosomal subunit | 1.097555 | 9.875817 |
| GO:0022627~cytosolic small ribosomal subunit | 1.126124 | 9.277004 |
| GO:0006446~regulation of translational initiation | 1.686112 | 4.77484 |
| GO:0003743~translation initiation factor activity | 3.064343 | 4.991915 |
| GO:0002181~cytoplasmic translation | 1.042256 | 3.180626 |
| GO:0001731~formation of translation preinitiation complex | 3.686284 | 3.049949 |
| GO:0006414~translational elongation | 3.817529 | 2.544042 |
| GO:0019843~rRNA binding | 2.895544 | 2.091965 |
| GO:0000470~maturation of LSU-rRNA | 3.817529 | 1.810927 |

##### Supplementary Table 2: Top ten GO terms enriched in RBN#3c CLIP peaks generated by DAVID analysis

| Category | Log2FoldChange | -log10FDR |
|---|---|---|
| GO:0046718~viral entry into host cell | -2.301341376 | 2.523992194 |
| GO:0001618~virus receptor activity | -2.350893537 | 2.523842019 |
| GO:0005903~brush border | -2.394218625 | 2.412101506 |
| GO:0015629~actin cytoskeleton | -1.450462735 | 2.184863887 |
| GO:0031965~nuclear membrane | -1.379443272 | 1.985604665 |
| GO:0044829~positive regulation by host of viral genome replication | -4.03830697 | 1.175787068 |
| GO:0000289~nuclear-transcribed mRNA poly(A) tail shortening | -2.765288476 | 1.11162171 |
| GO:0042555~MCM complex | -3.961259217 | 1.741727801 |
| GO:0008536~Ran GTPase binding | -2.698816841 | 1.236110328 |
| GO:0045862~positive regulation of proteolysis | -3.112307552 | 1.013102367 |

**Supplementary Table 3: List of long non-coding RNAs (lncRNAs) crosslinked to DEK in HeLa S3 cells**

| lncRNA name | Gene ID | Description |
|---|---|---|
| **CASC19** | 103021165 | Cancer Susceptibility Candidate 19; also named as cancer-associated region long non-coding RNA 6 (CARLos-6) in the 8q24 regions |
| **CCAT1** | 100507056 | Colon cancer-associated transcript-1 |
| **LINC00473** | 90632 | Regulate cAMP-mediated gene expression |
| **LOC100129434** | 100129434 | Uncharacterized feature |
| **NORAD** | 647979 | Necessary in genome stability |
| **UGDH-AS1** | 100885776 | UDP-glucose dehydrogenase |

**Supplementary Table 4: List of small nucleolar RNAs (snoRNAs) crosslinked to DEK in two individual DEK wt CLIP samples**

| snoRNA Name | snoRNA family | DEKwt_CLIP_1 | DEKwt_CLIP_2 |
|---|---|---|---|
| **SNORA73A** | box H/ACA | - | + |
| **SNORA73B** | box H/ACA | - | + |
| **SNORA10** | box H/ACA | + | + |
| **SNORD133** | Box C/D | + | + |
| **SNORD56B** | Box C/D | - | + |
| **SNORD3D** | Box C/D | - | + |
| **SNORD17** | Box C/D | - | + |
| **SNORD38A** | Box C/D | + | - |
| **RMRP** | MRP | - | + |

**Supplementary Table 5:** Oligonucleotides used for RIP-qPCR:

| Name | Sequence (5'→3') |
|---|---|
| **45S_F** | ACC CAC CCT CGG TGA GA |
| **45S_R** | AGT CGG GTT GCT TGG GAA TGC |
| **28S_F** | CCC AGT GCT TCT GAA TGT CAA |
| **28S_R** | CCC TTA CGG TAC TTG TTG ACT |
| **GAPDH_F** | GTC TCC TCT GAC TTC AAC AGC G |
| **GAPDH_R** | ACC ACC CTG TTG CTG TAG CCA A |
| **NORAD_F** | CTC TGC TGT GGC TGC CC |
| **NORAD_R** | GGG TGG GAA AGA GAG GTT CG |
| **Actin_F** | CTA TGC CTC TGG ACG CAC AAC T |
| **Actin_R** | CAG ATC CAG ACG CAT GAT GGC A |

**Supplementary Table 6:** Oligonucleotides used for Northern Blot/FISH:

| Name | Sequence (5'→3') | Labelling for Northern Blot | Labelling for FISH |
|---|---|---|---|
| **5'ITS1** | CCT CGC CCT CCG GGC TCC GTT AAT GAT C | 5'Biotin | 5'Cy5 |
| **ITS1-A** | AGG GGT CTT TAA ACC TCC GCG CCG GAA CGC GCT AGG TAC | 5'Biotin | N/A |
| **ITS1-B** | GAG TCC GCG GTG GAG | 5'Biotin | N/A |
| **ITS1-C** | GGT TGC CTC AGG CCG | 5'Biotin | N/A |
| **ITS2** | CTG CGA GGG AAC CCC CAG CCG CGCA | 5'Biotin | 5'Cy5 |
| **ITS2d.e** | GCG CGA CGG CGG ACG ACA CCG CGG CGT | 5'Biotin | N/A |
| **28S** | GAG GGA ACC AGC TAC TAG ATG GTT CGA TTA | 5'Biotin | 5'Cy5 |
| **18S** | TTT ACT TCC TCT AGA TAG TCA AGT TCG ACC | 5'Biotin | 5'Cy5 |

| | | | |
|---|---|---|---|
| **5S** | CCG AGA TCA GAC GAG ATC GGG CGC GTT CAG GGTGGT ATG G | 5'Biotin | N/A |
| **5.8S** | CAATGTGTCCTGCAATTCAC | 5'Biotin | N/A |

**Supplementary Table 7:** Oligonucleotides used for quantification of pre-rRNA species in total cell lysates:

| **Name** | **Sequence (5'→3')** |
|---|---|
| **ETS-1_F** | GTG CGT GTC AGG CGT TCT |
| **ETS-1_R** | GGG GAG AGG AGA GAC GAG |
| **H4_F** | CGA CGA CCC ATT CGA ACG TCT |
| **H4_R** | CTC TCC GGA ATC GAA CCC TGA |
| **ITS1_F** | CTC TCC GGA ATC GAA CCC TGA |
| **ITS1_R** | CGC GGA CAC CAC CCC ACA |
| **ITS2_F** | CCC GCC CCG CGG CCC GC |
| **ITS2_R** | CGA CGC GGA AGC TCG GGA |
| **H8_F** | AGT CGG GTT GCT TGG GAA TGC |
| **H8_R** | CCC TTA CGG TAC TTG TTG ACT |
| **DEK_F** | TGG GTC AGT TCA GTG GCT TTC C |
| **DEK_R** | CTC TCC AAA TCA AGA ACC TCA CAG |

Oligonucleotides used for HurDNA cloning in nuclear run-on assay (retrieved from [9]):

| **Name** | **Sequence (5'→3')** |
|---|---|
| **HurDNA_F** | CGC GGA TCC TGT CCT TGG GTT GAC CAG AG |
| **HurDNA_R** | CCG GAA TTC GCA AGT CGA CAA CCA CTG GA |
